# Genomic signatures of hybridization on the neo-X chromosome of *Rumex hastatulus*

**DOI:** 10.1101/2022.02.10.479988

**Authors:** Felix EG Beaudry, Joanna L Rifkin, Amanda L Peake, Deanna Kim, Madeline Jarvis-Cross, Spencer CH Barrett, Stephen I Wright

## Abstract

Natural hybrid zones provide opportunities for studies of the evolution of reproductive isolation in wild populations. Although several recent investigations have found that the formation of neo-sex chromosomes is associated with reproductive isolation, the mechanisms remain unclear in most cases. Here, we assess the contemporary structure of gene flow in the contact zone between largely allopatric cytotypes of the dioecious plant *Rumex hastatulus*, a species in which there is evidence of sex chromosome turn-over. Males to the west of the Mississippi river, USA, have an X and a single Y chromosome, whereas populations to the east of the river have undergone a chromosomal rearrangement giving rise to a larger X and two Y chromosomes. Using reduced-representation sequencing, we provide evidence that hybrids form readily and survive multiple backcross generations in the field, demonstrating the potential for ongoing gene flow between the cytotypes. At the scale of chromosomes, cline analysis of each chromosome separately captured no signals of difference in cline shape between chromosomes. However, when comparing SNPs, principal component regression revealed a significant increase in the contribution of individual SNPs to inter-cytotype differentiation on the neo-X, but no correlation with recombination rate. Cline analysis revealed that the only SNPs with significantly shallower clines than the genome-average were located on the neo-X. Our data are consistent with a role for the neo-sex chromosome in reproductive isolation between *R. hastatulus cytotypes*. Our investigation highlights the importance of studying plant hybrid zones in species with sex chromosomes for understanding mechanisms of reproductive isolation and for understanding the role of gene flow in governing the spread of the neo-X chromosomes.

## INTRODUCTION

Where species meet and breed successfully in the wild, hybrid zones can form. In these zones of admixture, segregation and recombination shuffle the genomes of closely related species (Barton and Hewitt, 1985; Gompert et al. 2017; Harrison 1993). Allele frequencies often display a geographic pattern of greater admixture near the center of the hybrid zone, called an allele frequency cline (Barton, 1979; Hewitt, 1989; Rieseberg and Wendel 1993). Under continuous gene flow, hybrid genomes are expected to contain a mixture of alleles from both species at all sites (Cruzan and Hendrickson 2020; Harrison 1993; Hewitt 1988). However, allele frequency clines are often heterogeneous across the genome of hybrid individuals (Aeschbacher et al. 2017; Brandvain et al. 2014; Gibson and Moyle 2019; Rieseberg et al 1995; Rifkin et al. 2019). Genomic regions resistant to the shuffling of alleles can maintain reproductive incompatibilities between species, and loci at which the allele frequency cline is steep can be an indication of genes involved with such incompatibilities (Barton 1983; Barton and Bengtsson 1986; Cruzan et al 2021; Gompert et al. 2017). Hybrid zones can therefore provide opportunities for direct insights into the genetic mechanisms governing reproductive isolation in natural populations.

Chromosomal rearrangements between closely related species, including translocations and inversions, often appear as islands of increased genetic divergence with steeper allele frequency change across hybrid zones compared with genome averages (Coughlan and Willis 2019; Lee et al. 2017; Noor et al. 2001; Todesco et al. 2019). Rearrangements are thought to be involved in hybrid break down because of their impact on the pairing of homologous chromosomes at meiosis. In early hybrid generations, failures in pairing can lead to hybrid offspring with missing or extra chromosomes, with strongly deleterious effects (Federley 1913; Torgasheva and Boroding 2016). In addition, failures in pairing of homologous chromosomes may play a role in restricting recombination locally in the genome without causing hybrid failure (Rieseberg 2001). Recombination rate is generally thought to be an important driver of speciation because it can keep alleles involved in adaptation together when recombination rate is low (Todesco et al. 2019), or alternatively purge deleterious incompatibilities when recombination rate is high (Schumer et al. 2018).

Sex chromosomes are disproportionately associated with genomic divergence and reproductive isolation (Coyne and Orr 1989; Haldane 1922; Presgraves 2018), and the X chromosome is frequently found to show steeper clines across hybrid zones than other genomic regions (Geraldes et al. 2006; Hooper et al. 2018; Moran et al. 2018; Tucker et al. 1992). Sex chromosomes may contribute to species incompatibilities when recessive X alleles are unmasked by Y chromosome degeneration, or if genes with sex-specific fitness effects tend to evolve incompatibilities faster than other genes and disproportionately accumulate on the X (Lasne et al. 2017; Wu 2001). Growing empirical evidence supports sex chromosome rearrangements, such as the formation of neo-sex chromosomes, as being associated with the speciation process in animals (Benatti et al. 2010; Bracewell et al. 2017; Franchini et al. 2018; Kitano et al. 2009; Smith et al. 2016; Wang et al. 2021) and plants (Beaudry et al 2019). But the mechanisms underlying their association with reproductive isolation remain unclear, especially in plants. Studies of hybrids between species that differ in sex chromosome rearrangement are of particular interest for disentangling the intrinsic factors governing the genetics of reproductive isolation.

In the dioecious plant *Rumex hastatulus* (Polygonaceae), males to the west of the Mississippi river USA have an X and a single Y chromosome (hereafter XX/XY cytotype), whereas populations to the east of the river have a larger X and two Y chromosomes (hereafter XX/XY_1_Y_2_ cytotype). The second Y in the XX/XY_1_Y_2_ cytotype arose from a chromosomal translocation between the ancestral X and an autosome (Autosome 3), leaving the autosomal homolog of the translocated X to segregate as a second Y chromosome and resulting in XYY males (Kasjaniuk et al. 2018; Rifkin et al. 2021; Smith 1963). Hereafter, we refer to the newly sex-linked region as the ‘neo-X’ and the chromosome that incorporates both the ancestral- and neo-X as the ‘fused-X’. The role of neo-sex chromosomes in reproductive isolation and the consequences for the hybridization process in the contact zone of *R. hastatulus* cytotypes remain unresolved.

The first evidence that the different geographic cytotypes of *R. hastatulus* are in the early stages of reproductive isolation came from experimental crosses between individuals of different cytotypes. This research revealed that F_1_ hybrid pollen had reduced viability, likely associated with uneven segregation of the sex chromosomes in hybrids (Kasjaniuk et al. 2018). Additional evidence that the neo-sex chromosome plays a key role in reproductive isolation comes from demographic inference from allopatric populations where a temporal association between the sex chromosome rearrangement and a loss of gene flow between the two cytotypes was observed (Beaudry et al. 2019). However, earlier cytotype analysis reported a substantial zone of hybridization around the Mississippi river region (Jackson, 1967; Smith 1969). But whether gene flow in the zone of contact is ongoing past the *F*_1_ generation remained unclear thus motivating our study.

Genomic studies in contact zones can provide insight into the prevalence of hybrids, the viability of hybrid offspring, and the degree of heterogeneity in gene flow across the genome (Abbott 2017; Barton and Hewitt 1985; Gompert et al. 2017). Although investigations of this type can provide critical insights into the process of speciation, hybrid zones of plant species with sex chromosomes are generally under-investigated at the genomic level (Pickup et al. 2019). Here, we assessed the contemporary structure of gene flow between the cytotypes of *R. hastatulus* by analysing individuals in the contact zone originally identified by Smith (1969). Although previous work in allopatric populations of the cytotypes focused on demographic inference from population genetic summary statistics (Beaudry et al. 2019), here we directly investigated the propensity of individuals to hybridize and the outcomes of such hybridization in contact zone populations. The primary goal of our study was therefore to establish the genomic structure of gene flow across the *R. hastatulus* cytotype contact zone and in so doing address the following questions: 1) Is gene flow occurring and do hybrids form? 2) If so, what is the location and breadth of the hybrid zone? 3) What are the relative roles of recombination, chromosomal rearrangements, and sex-linkage on the extent of gene flow in the hybrid zone?

## METHODS

### Population-level sampling

As cytotype hybrids are not reliably distinguished by observable phenotypic traits (Jackson 1967; Smith 1969), we used genetic data to quantify the extent of hybridization in the zone of contact between the cytotypes by collecting samples from the hybrid zone. To identify the hybrid zone, we mapped populations of hybrids identified by Smith (1969; Fig. 1 therein) to GPS coordinates using QGIS (grey in Fig. 1 here). In June 2018, we collected seed from a minimum of 20 open-pollinated female plants in each of 30 populations approximately equally distributed among three transects positioned across both sides of the Mississippi river in the US states of Mississippi, Louisiana, and Arkansas (yellow circles in Fig. 1, S1). To maximize the likelihood of encountering the zone of contact and collecting hybrids, sampling transects spanned both sides of the Mississippi river. Although individuals were not karyotyped, we assumed that plants at the extreme opposite ends of the range we sampled are individuals from the different cytotypes because longitude is a strong predictor of karyotype identity (Smith 1963), an assumption supported by linkage group analysis (see Rifkin et al. 2020). Consistent with data from online herbaria (Data Portal 2022), we observed a notable discontinuity in *R. hastatulus* occurrence close to the Mississippi river at the transition into Major Land Resource Area (MLRA) region O: *Mississippi Delta Cotton and Feed Grains Region* (Fig. S1).

**Figure 1.**
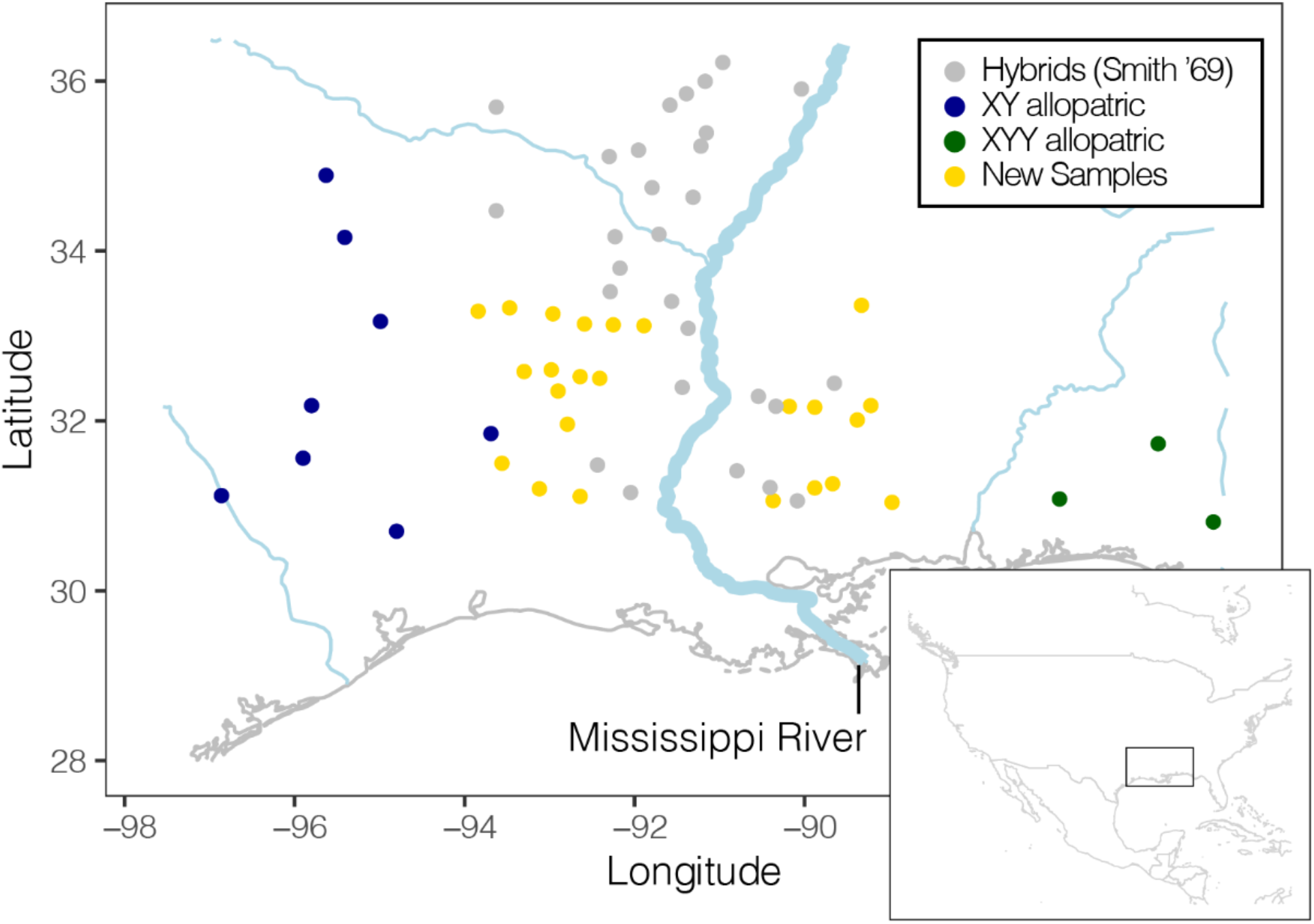
Samples of *Rumex hastatulus* across the hybrid zone that were sequenced: new samples (“contact zone”) in yellow, subset of allopatric XY (blue) and XYY (green) samples previously sequenced in Beaudry et al. (2019), and samples identified as hybrids in Smith (1969) in grey which were not sequenced. Coastline as grey line, rivers as blue lines; Mississippi river is bolded and labelled. Inset: Map of North America with black box showing region sampled.

For subsequent genomic analysis, we sampled 20 seeds from each of 20 maternal families among the 30 populations (12,000 seeds from 600 maternal families in total; see Fig. S2 for experimental design). Seeds were bleached, rinsed, chilled at 4°C, and germinated on moist filter paper in growth chambers for two weeks. We transferred germinated seeds to 10cm pots following earlier protocols and these were grown until they began flowering in the glasshouse (see Pickup et al. 2013 for details). One individual from each maternal family was chosen for DNA extraction, to a maximum of 10 individuals from different families per population. As not all individuals survived to first flower, this resulted in 143 leaf tissue samples among 24 populations (see Table S1 for population-level survival information). In six populations, all individuals died before flowering (X-ed yellow circles in Fig. S1). For individuals that survived until flowering, we sexed samples using floral phenotype (presence of anthers vs stigma) and collected tissue from young leaves for DNA extraction. As previous work anecdotally recorded a low level of mismatch between sex phenotype and sex genotype for some samples (Beaudry et al. 2019), we froze flower samples to confirm the match between genotypic and phenotypic sex. Out of 143 genotyped samples, 4 individuals had a mismatch between phenotype and genotype (the presence/absence of diagnostic X-Y fixed differences shared by both allopatric populations); these individuals were recorded as having inconclusive sex and were excluded from analyses considering sex chromosomes.

### DNA extractions and sequencing

We extracted DNA using a DNeasy Plant Mini Kit (QIAGEN GmbH, Hilden, GERMANY) following the manufacturer’s protocol. The Elshire group in Palmerston, NZ performed library preparation of the DNA samples using the genotype-by-sequencing (GBS) method (Elshire et al. 2011); briefly, reads were fragmented using the restriction enzyme PstI and ligated to barcodes. Libraries were sequenced over 1 lane of 2 × 150bp Illumina HiSeq at the Australian Genome Research Facility (AGRF) in Westmead, Australia. Sequenced libraries were demultiplexed with axe-demux (Murray and Borevitz 2018). Barcodes and adapters were removed using trimgalore v.0.4.3 (https://www.bioinformatics.babraham.ac.uk/projects/trim_galore/) according to default parameters.

### SNP calling

We combined our sequencing data with 94 GBS libraries from 21 allopatric populations (Fig. 1, Fig. S1) previously sequenced in Beaudry et al. (2019). Hereafter, the samples sequenced in this study are referred to as ‘contact zone’ whereas those from Beaudry et al. (2019) are referred to ‘allopatric’. We aligned sequencing reads from all 237 samples (94 allopatric and 143 contact zone individuals) from 45 populations (21 allopatric and 24 contact zone populations) in our dataset to the repeat-masked *R. hastatulus* XX/XY-cytotype genome assembly; see Rifkin et al. (2020) for genome assembly and repeat-masking methods. We performed alignments using default parameters for paired reads in NextGenMap v.0.5.5 (Sedlazeck et al. 2013). Bam files were sorted and read groups added with Picard v.2.23.8 (http://broadinstitute.github.io/picard). After filtering individuals based on a 5x minimum mean genome-wide read coverage cutoff, we retained 140 contact zone samples and all 94 allopatric samples (for 234 samples total). Variant sites were called using the commands *mpileup* followed by *call* in bcftools v. 1.9-67-g626e46b (Li 2011) applying no filtering at this stage. We filtered the resulting .vcf using vcftools v. 0.1.15 (Danecek et al. 2011) for biallelic SNPs with sequencing information for more than 60% of individuals, a minimum PHRED quality score of 10, an all-sample mean site depth of greater than 5 reads, a minimum genotype PHRED quality score of 5 and a minor allele frequency (MAF) of greater than 1% (i.e., alleles present in only 2 copies or less were removed). We also removed all SNPs where all individuals were heterozygous, or where all individuals from any one population were missing data. This filtering yielded 13,578 SNPs.

### Removing X-Y fixed differences

Because of the divergence of the Y chromosome from the X chromosome and of the neo-Y from the neo-X (see Hough et al. 2014), some loci identified as SNPs may be fixed differences between X and Y (or neo-X and neo-Y). This would be misleading for inferences of hybridization because these loci inherently show high heterozygosity in males, as well as strong frequency differences between the sexes. To remove X-Y fixed differences, we used differences in heterozygosity between males and females by calculating expected heterozygosity (*2pq*) in all males or all females, respectively, in each of four categories (from west to east): allopatric XX/XY, western contact zone, eastern contact zone and allopatric XX/XY_1_Y_2_. We then subtracted male heterozygosity from female heterozygosity to estimate excess female heterozygosity on autosomes, the X and neo-X chromosomes (Fig. S3). Taking the set of SNPs with male excess heterozygosity >0.25 (−0.25 in Fig. S3) as potentially capturing Y-specific polymorphism and fixations, we removed 263 SNPs yielding a total of 13,315 SNPs for our primary analysis. The one part of our study where these SNPs are not removed is the analysis of clines SNP-by-SNP (using *bgc*, see below) in order to avoid excluding highly differentiated X-linked SNPs.

### Individual-level genotype-based hybrid assessment

We first performed a Principal Component Analysis (PCA) on individual genotypes (coded as [0,1,2]) using FactoMineR (Husson et al. 2010) in R. We next used the probabilistic model analysis software STRUCTURE v. 2.3.4 (Pritchard et al. 2000), with clusters (K) ranging from 2 to 5, and ran six MCMC chains for a 1,000 generation burn-in followed by 10,000 generations of analysis (11,000 iterations total) for each value of K. MCMC chains for STRUCTURE runs K>3 failed to converge and were not analysed further (CONSTRUCT analysis suggests K>3 is not representative of further population substructure, see below). We aligned STRUCTURE runs by clusters using permutations in CLUMPP v.1.1.2 (Jakobsson and Rosenberg 2007). To investigate population substructure specific to the XX/XY cytotype, we used PCA only on individuals from these populations. We estimated linkage disequilibrium using plink flags --r2 inter-chr -and --ld-window-r2 0.

We examined the relations between heterozygosity and ancestry, using graphical approaches derived from the de Finetti diagram (Cannings and Edwards, 1968), to infer the generation at which hybrids may be inviable or infertile and, thus, to assess the genetic characteristics of hybrid incompatibilities (see Turelli and Orr 2000). We jointly inferred Hybrid Index (HI) and heterozygosity using HIEST v. 2.0 (Fitzpatrick 2012). For a set of SNPs for which alleles were diagnostic to each population, first-generation hybrids are expected to show peak heterozygosity and a 50/50 ancestry mix, whereas individuals from populations without gene flow are expected to show both low heterozygosity and exclusive ancestry. Because HIEST assumes the reference allele in the genome assembly is diagnostic for the reference population, we assumed the reference base, which is an XX/XY-cytotype male (Rifkin et al. 2020), is representative of the XX/XY-cytotype as a whole. We began our analysis using all SNPs, but these results did not conclusively distinguish between the cytotypes, likely due to the predominance of shared SNPs between cytotypes and low-frequency unique SNPs (Beaudry et al. 2019). To increase our resolution to detect signatures of hybridization, we repeated the analysis using only the 100 SNPs with the largest contributions to PC1.

### Population-level allele-frequency-based hybrid assessment

Using the same method as outlined in the preceding section but on population-level allele frequencies, we performed both PCA and assessed population-level clustering using CONSTRUCT v1.0.4 (Bradburd et al. 2018) in R. We ran both spatial and non-spatial runs of CONSTRUCT for K-values between 1-6 and visualized CONSTRUCT model comparisons using the calculate.layer.contribution function. To investigate rates of effective migration across the landscape in two-dimensions, we also evaluated the Estimate Effective Migration Surface v.0.0.0.9 (EEMS; Petkova et al. 2017). As our data included some missing genotypes, we converted the .bed file to .diffs using bed2diffs_v2 in EEMS. We ran six MCMC chains for 2,000,000 iterations with identical parameters among chains with a burn-in length of 1,000,000 iterations, a thinning interval of 9999 iterations, and a model with 800 demes.

### Cline fitting

As canonical cline-based analyses depend on diagnostic fixed differences between groups, the absence of fixed differences in our genetic markers made the classic approach to cline analysis intractable. Instead, by assuming Hybrid Index is continuous across the hybrid zone, we fitted allele frequency clines to Hybrid Index values, allowing us to estimate the expected selection against hybrids at randomly chosen loci in the genome (Gompert et al. 2017; Hooper et al. 2018). Because CONSTRUCT incorporates uncertainty of parental genotypes, it is a flexible tool to estimate Hybrid Index (Bradburd et al. 2018; Pritchard et al. 2000). For our Hybrid Index estimate, we used K=2 and a non-spatial model. To increase our sampling density of the putative hybrid zone, we combined data from all three transects. We confirmed our results with cline fitting to normalized population-level (allele-frequency) PC1 (*i*.*e*., using the same data as in CONSTRUCT) to ensure that our results were not influenced by over-fitting (see Wang et al. 2019).

Using HZAR (Derryberry et al. 2014), we fitted six cline models (plus a null model) for *p*, our standardized values of PC1 or HI estimated by (non-spatial) CONSTRUCT (see Table S2 for model summaries). Our null model (model 0) was that longitude explained no change in allele frequency, *i*.*e*., *p* is best explained as a horizontal line across the hybrid zone. We fitted two sets of models that allowed *p* to increase with longitude, the first of which had *p* starting at 0 and increasing to 1 (model 1-3) and a second that fitted a minimum (*p*_min_) and maximum (*p*_max_) value to *p*, according to the data (model 4-6). Models 1 and 4 followed a logit function, as would be expected if allele frequency change can be explained by dispersal alone. If selection is acting, change in *p* across longitude should depart from a logit function according to the parameters **δ**, the distance from the cline center where exponential decay in *p* starts, and ***τ***, the slope of linear decay in *p*. For our models including **δ** and ***τ***, we fitted a model where both **δ** and ***τ*** take the same value at each end of the hybrid zone (mirrored; models 2 and 5), whereas a second model set allowed for separate values of **δ** and ***τ*** on each side of the cline (models 3 and 6). Because we fitted models to both the PCA and CONSTRUCT values, this resulted in a total of 14 models. We used the convention of a two-point AIC difference for significance (Gay et al. 2008; Raufaste et al. 2005). We then repeated the above analyses on each chromosome, splitting each chromosome into a region of low (<0.1 cM/mb) or high (≥0.1 cM/mb) recombination, separately (see below for methods for recombination rate estimation).

### Genomic heterogeneity in hybridization

We evaluated the effect of recombination density [recombination rate (cM) per megabase (mb)] on differentiation between the cytotypes by estimating the recombination intensity (cM/mb) in 1mb windows along the genome using the anchored linkage map from Rifkin et al. (2020). To estimate recombination rate, we fitted polynomials onto the anchored linkage map, using the method and adapted code from Corbett-Detig et al. (2015).

We estimated heterogeneity in clines SNP-by-SNP using Bayesian genomic cline *bgc* (Gompert and Buerkle 2011). In contrast to geographic cline analysis, genomic cline analysis is based on identifying loci with exceptional allele frequency differences between hybrids and parental populations, given a base level of introgression also estimated by the model. Rather than explicitly estimating change in allele frequency across space, *bgc* fits allele frequency clines to every SNP. In *bgc*, the cline is designated by two parameters: **α** represents the probability of an allele in the hybrid population coming disproportionately from one parental population and **β** represents the rate in transition from one parental population to the other (see Gompert and Buerkle 2011). We converted SNP files from vcf to *bgc* format using the vcf2bgc.pl script in *bgc* v.1.0. This was then run for 50,000 MCMC steps (-x), discarding the first 10,000 steps as burnin (-n) and thinned for every 5th step (-t) using the genotype uncertainty model (-N) and including precision parameters (-p) and quantiles (-q). We returned estimates from all MCMC runs using the estpost v 1.0 with credible interval set to default (*i*.*e*., 0.95).

## RESULTS

### Genomic evidence for hybrid individuals

To identify the occurrence of hybrid individuals in the zone of contact between the *R. hastatulus* cytotypes, we first performed genetic clustering analyses using 13,315 SNPs for 234 individuals from 45 populations (21 allopatric and 24 contact zone; Fig. 1, S1). We began by performing PCA, and found that PC1 grouped all individuals from allopatric populations into two separate clusters (empty circles in Fig. 2A). PC1 was strongly correlated with longitude (Pearson Product-Moment Correlation: 0.924; *t* = 36.83, D.F. = 232, *P* < 2.2e-16). Individuals from a region of about 200km on the eastern side of the Mississippi river were continuously distributed between the two cytotypes (full circles in Fig. 2B). The occurrence of continuous variation in genotype-space in the geographic region between the allopatric populations is consistent with the hypothesis that hybrids occur in this range. Hereafter, we therefore refer to this geographic region as the hybrid zone. Most of the variance in PC2 was restricted to individuals from the hybrid zone (Fig. 2A), consistent with simulation work showing hybrid populations tend to harbor unique genotypic variation (Gompert and Buerkle 2016).

**Figure 2.**
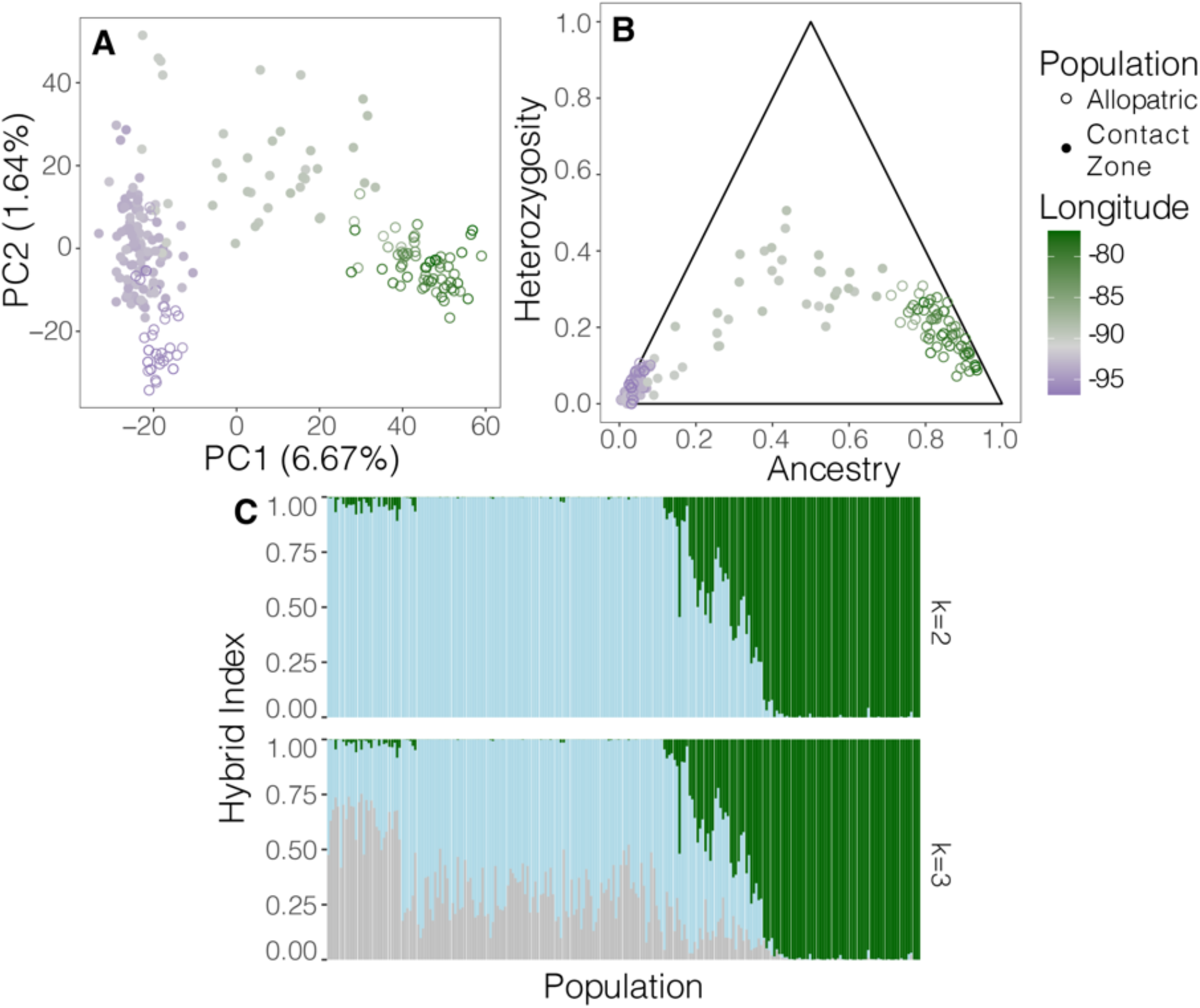
Inferring hybridization between cytotypes of *Rumex hastatulus*. **A**. Principal component analysis. **B**. Hybrid generation assessment for top 100 SNPs of PC1 by assessing estimated ancestry compared to heterozygosity. In both plots, individuals are colored by the longitude at which they were sampled [purple longitude = 100° W, gray longitude = 90° W (Mississippi River), green longitude 75° W.] and shape represents whether individuals were collected from the contact zone or allopatric populations. **C**. Individual-based STRUCTURE analysis, with populations of *R. hastatulus* sorted west (left) to east (right). White vertical lines are breaks between populations.

The results of our STRUCTURE analysis (assuming two clusters i.e. K=2) separated individuals into geographic groups corresponding to the inferred division between the cytotypes (Fig. 2C). Individuals from the hybrid zone exhibited Hybrid Index (HI) values between 0.04 and 0.75, whereas all individuals from allopatric populations had scores higher than 0.95 or lower than 0.05. With K=3, the data suggest geographic substructure specific to the XX/XY cytotype, discussed further below. Hybrid Index values were highly correlated with PC1 (Pearson Product-Moment Correlation: 0.989; *t* = 101.26, D.F. = 234, *P* < 2.2e-16; Fig. S4A).

We next jointly inferred hybrid index (ancestry) and heterozygosity to assess the number of generations for which hybridization has successfully proceeded. Using all SNPs, the two cytotypes were not conclusively distinguished (Fig. S5). To increase our resolution to detect signatures of hybridization, we repeated the analysis using only the 100 SNPs with the largest contributions to PC1 (Fig. 2B). In this second analysis, allopatric populations tended towards the corners of the triangle, whereas most individuals from populations in the hybrid zone were in the center of the triangle. Our results do not conclusively separate individuals into generations; however, the continuous patterns of ancestry and heterozygosity (Figure 2) are consistent with late-generation hybrids.

### Gene flow across the hybrid zone

In contrast to the individual-level genotype-based analyses of the previous section, we next conducted analyses on population-level allele frequencies. PCA of allele frequencies replicated the individual level analyses: PC1 was strongly correlated with longitude (Pearson Product-Moment Correlation: 0.925; *t* = 15.98, D.F. = 43, *P* < 2.2e-16; Fig. S6). Population-level clustering analysis (using CONSTRUCT; Bradburd et al. 2018) separated the cytotypes into separate clusters (Fig. S7). Non-spatial CONSTRUCT model comparisons suggested three clusters (K=3) partitioned by the largest portion of genotype variance (Fig. S7A), whereas spatial runs suggested increased values of K do not explain more genotype variance. CONSTRUCT at K=3 displayed a pattern concordant with the STRUCTURE results, splitting by cytotype and suggesting population substructure specific to the XX/XY-cytotype in both spatial and non-spatial runs (Fig. S7D, B). Regardless of K or whether the model was spatial or not, CONSTRUCT results confirmed that the majority of populations in the 200km region east of the Mississippi river are composed of hybrid individuals. Calculated at the population level, Hybrid Index was correlated with PC1 using population allele frequency data (Pearson Product-Moment Correlation: 0.996; t = 77.38, D.F. = 43, *P* < 2.2e-16; Fig. S4B).

We next fitted allele frequency clines for seven models for normalized allele-frequency PC1 and the Hybrid Index estimated by CONSTRUCT assuming a non-spatial model and K=2 (Fig 3; Table S2 for model summaries). Briefly, parameters were sequentially added to the cline equation, starting from a null model, to include two slope parameters (center and width; model 1), two decay parameters (**δ**, the distance from the cline center where exponential decay in *p* starts, and ***τ***, the slope of linear decay in allele frequency; model 2), independent decay parameters for each allopatric population (model 3) and an allele frequency in allopatric population parameter (rather than assume alleles were alternately fixed; model 4-6. For CONSTRUCT, model 2 (slope term and decay term; AICc=24.03) was marginally more likely than 4 (slope terms [center and width], and allopatric populations unfixed; AICc=24.20), but both models were more significant than any other model (Table 1; purple in Fig 3). For normalized PC1, model 4 (AICc=17.59) was marginally better supported than model 2 (AIC=17.65) and both models were significantly more supported than other models (Table 1; red in Fig 3). Whereas model 2 suggests fixed differences but a decay term, model 4 suggests no decay term but differences between allopatric populations are not fixed. Based on the 95% confidence intervals, clines fit to both CONSTRUCT and PCA were concordant for both cline center and width (Table 1; Fig. S9).

**Figure 3.**
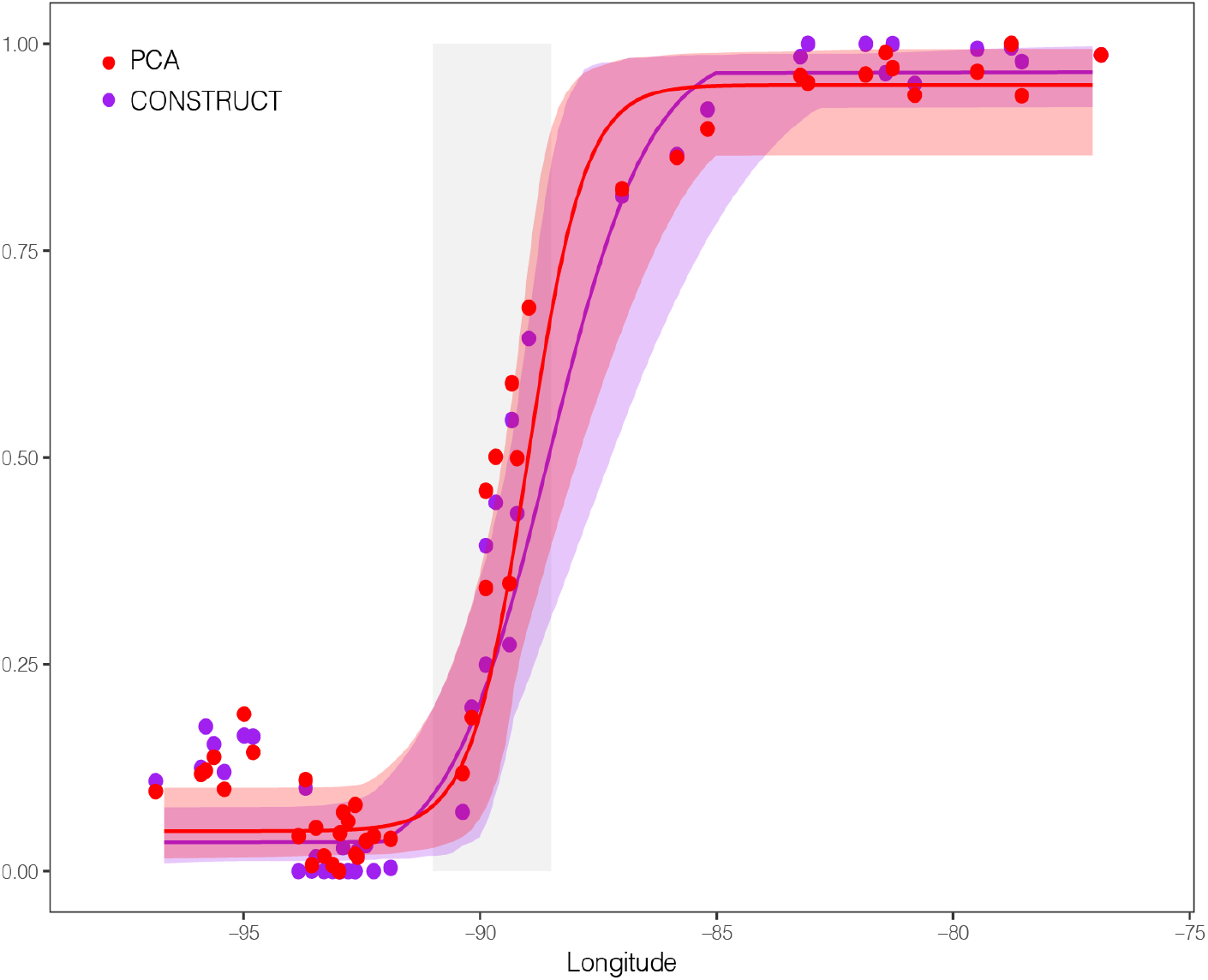
The rate of change in hybrid index across longitude for population samples of *Rumex hastatulus*. Points are sampling locations, black line is the best fit estimated by HZAR and the gray shaded ribbon is the confidence interval. Blue = PC1, Red = CONSTRUCT.

**Table 1.** Results of cline fitting models with HZAR using both normalized PC1 and Hybrid Index (H.I.) estimated at the population-level by CONSTRUCT in *Rumex hastatulus*.

| Cline | Model | AICc | Parameter | Estimate | Confidence interval |
| --- | --- | --- | --- | --- | --- |
| CONSTRUCT H.I. | 2 | 24.03 | Center (Lon.) | -88.55° | [-89.30; -87.53] |
|  |  |  | Width (km) | 385.79 | [122.45; 615.94] |
| | | | $\delta$ (km) | 318.40 | [84.37; 505.38] |
| | | | $\tau$ | 0.0041 | [7.41e-06; 0.34] |
| Norm. PC1 | 4 | 17.59 | Center (Lon.) | -89.01° | [-89.47; -88.03] |
|  |  |  | Width (km) | 212.06 | [93.80; 492.84] |
| | | | $p_{\min}$ | 0.05 | [0.019; 0.095] |
| | | | $p_{\max}$ | 0.95 | [0.88; 0.99] |

The estimated effective migration surface (EEMS; Petkova et al. 2017) revealed significant signatures of reduced gene flow across the Mississippi river (Posterior Likelihood of log(p) > 0, *P* < 0.05), with lower rates of gene flow between populations on the east side of the river compared to the west side (Fig. S8). While gene flow was prevalent between populations on the west side of the river (characterized as having low hybrid index values), gene flow was restricted between populations on the east side of the river, where we identified the hybrid zone. This result suggests that gene flow is lower across the hybrid zone than in other geographic areas that we sampled.

### Genomic heterogeneity in hybridization

We next aimed to disentangle the relative roles of sex-linkage and recombination in hybridization between the XX/XY and XX/XY_1_Y_2_ cytotypes. First, we ran CONSTRUCT followed by cline fitting of Hybrid Index for each chromosome separately and within each chromosome we split the chromosome into low (cM/mb < 0.1) and high (cM/mb ≥ 0.1) recombination regions (Table S3 and Fig S10). The estimated center of the cline was consistent across chromosomes, between -88.51° and -89.06° (east of the Mississippi river), and also consistent with the genome-wide average. Because all confidence intervals overlapped, we concluded that there was no significant difference between our parameter estimates for either cline width or cline center position.

We assessed the relative contribution of SNPs to genotype (individual-level) PC1. To this end, we calculated the average effect size (loading) of SNPs on PC1 in windows across the genome. To jointly evaluate the impact of recombination intensity, we fitted a linear model to calculate the effect of chromosome and recombination rate on PC1 effect size. We found the neo-X/A3 had a significantly increased contribution to PC1 effect size (Estimate = +0.0065, s.d.= 0.0015, *t*-value = 4.46, *P =* 9.98e-06), as did Autosome 4 (Estimate = +0.0043, s.d.= 0.0011, *t*-value = 4.04, *P =* 6.07e-05) compared to other chromosomes. However, we found no significant impact of recombination rate on PC1 loading (*P =* 0.45).

We next estimated heterogeneity in clines SNP-by-SNP. First, we evaluated broader patterns of male-female heterozygosity across chromosomes. Excess heterozygosity in males compared to females (< -0.3) likely represents polymorphism on the X or Y, whereas - 0.5 should represent fixed X-Y differences. Excess heterozygosity in females likely represents genes/sites missing from the Y (*i*.*e*. males are hemizygous at those loci). In three locations (XX/XY allopatric, western contact zone, and eastern contact zone), we observed more heterozygote excess on the X than on autosomes for both males and females, whereas we found a paucity of female heterozygote excess in allopatric XX/XY_1_Y_2_. In both allopatric and western contact zone populations, neo-X and its autosomal homolog in the XY-cytotype (Autosome 3) had identical patterns of heterozygosity as the autosomes, as would be expected because these sites are segregating autosomally in these locations. In the hybrid zone, females exhibited an excess of heterozygosity compared to males on the neo-X/A3 and above levels for the autosomal loci. In contrast, male-biased heterozygosity was not substantially different from the autosomal patterns on the neo-X/A3 in the hybrid zone. In allopatric XX/XY_1_Y_2_, we observed the highest increase in male-biased heterozygosity on the neo-X/A3. Given this latter pattern, high male excess heterozygosity may reflect greater levels of neo-X differentiation between the cytotypes.

We assessed clines SNP-by-SNP with Bayesian genomic cline (bgc) analysis, including our set of SNPs with excess male heterozygosity, as these may be due to high cytotype divergence on the neo-X. The direction of gene flow (**α**) was on average positive (estimate = 0.0163) and significantly different from zero (one sample t-test, *t* = 3.2329, d.f. = 13314, *P* = 0.001228), indicating a disproportionate representation of alleles from the XX/XY cytotype in hybrids probably as a result of asymmetric gene flow from the XX/XY cytotype. Outliers of **α** were evenly distributed across the genome. We detected three SNPs with positive outlier cline steepness values (**β**). These SNPs were all sex-linked, either on the X or neo-X (Fig 4A). Investigation of genotype frequencies in males and females supported that these SNPs are linked to the X or neo-X rather than the Y or neo-Y, given the pattern of homozygous genotype differences in females across the hybrid zone (Fig. 4B). In a linear model evaluating the impacts of sex-linkage and chromosome on **β**, sex-linked X was significantly higher than the genome average (Estimate =0.0152215, s.d.=0.0052071, *t*-value =2.923, *P =*0.00347), the PAR of the neo-X was significantly higher (Estimate =0.0148369, s.d.=0.0024659, *t*-value =6.017, *P =*1.84e-09) and the sex-linked portion of the neo-X had the largest effect (Estimate =0.0462763, s.d.=0.0049767, *t*-value =9.299, *P* < 2e-16; Fig. 4A). We identified a pair of SNPs 10bp apart with negative outlier **β** values (cline shallowness) on autosome 2.

**Figure 4.**
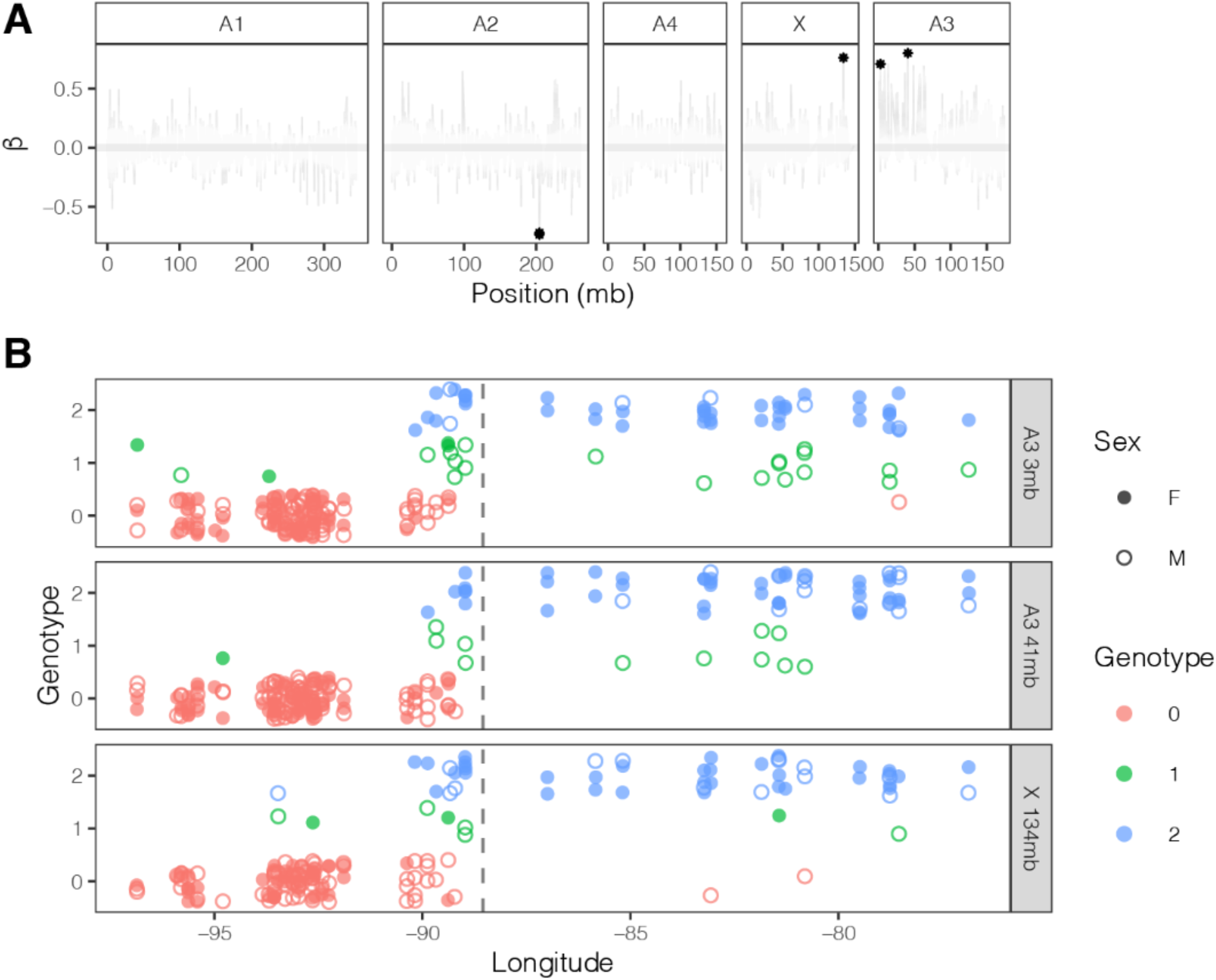
**A β** values across *Rumex hastatulus* chromosomes. Positive values indicate narrow clines (less introgression than expected; selection against hybrids) and negative values indicate shallow clines (more introgression than expected). The stars are the two outliers. **B** Genotypes for three outlier positive **β** SNPs across longitude. Circles represent sex (filled female, empty male) and color and y-axis are the same data for genotype. The dashed line is the best fit of the cline center.

### XX/XY cytotype substructure

Our STRUCTURE analysis suggested substructure in the XX/XY cytotype and therefore we performed PCA on genotypes specific to samples from this cytotype. Because the hybrid zone is located exclusively on the east side of the Mississippi, we included only individuals from the west side in this analysis. XX/XY-exclusive PC1 correlates with PC1 from the all-samples PCA (Pearson’s product-moment correlation = 0.57, *t* = 7.96, D.F. = 131, *P* = 7.24e-13), indicating we are identifying the same pattern in both PCAs. Also, it is evident that the XX/XY-exclusive PC1 correlates with longitude (0.70, *t* = 11.34, D.F. = 131, *P* < 2.2e-16). To locate the genomic source of XX/XY substructure, we investigated the effect size of SNPs on PC1 of the XX/XY-exclusive PCA. We found a sizeable region with many SNPs of large effect on the left end of autosome 2 (Fig. 5A). The region on the left is noteworthy for containing two nested inversions, based on comparative linkage mapping from mapping populations of the two cytotypes (Fig. 5B, data from Rifkin et al. 2020). Furthermore, based on our SNP set, we found high levels of linkage disequilibrium around the region of the inversion on A2 (Fig. 5C). A similar pattern appears for lower principal components: SNPs that contribute most to PCs 2-4 also align with regions where LD is disproportionately high. Rifkin et al. (2020) identified inversions between the cytotypes; the noteworthy exception is the peak in the pericentromeric region of Autosome 1 (A1). Although we do not have evidence that the inversions that appear in our comparative maps are segregating in the XX/XY-cytotype, the signal of population substructure specific to the XX/XY cytotype aligns well with these inversions and the regions of high linkage disequilibrium.

**Figure 5.**
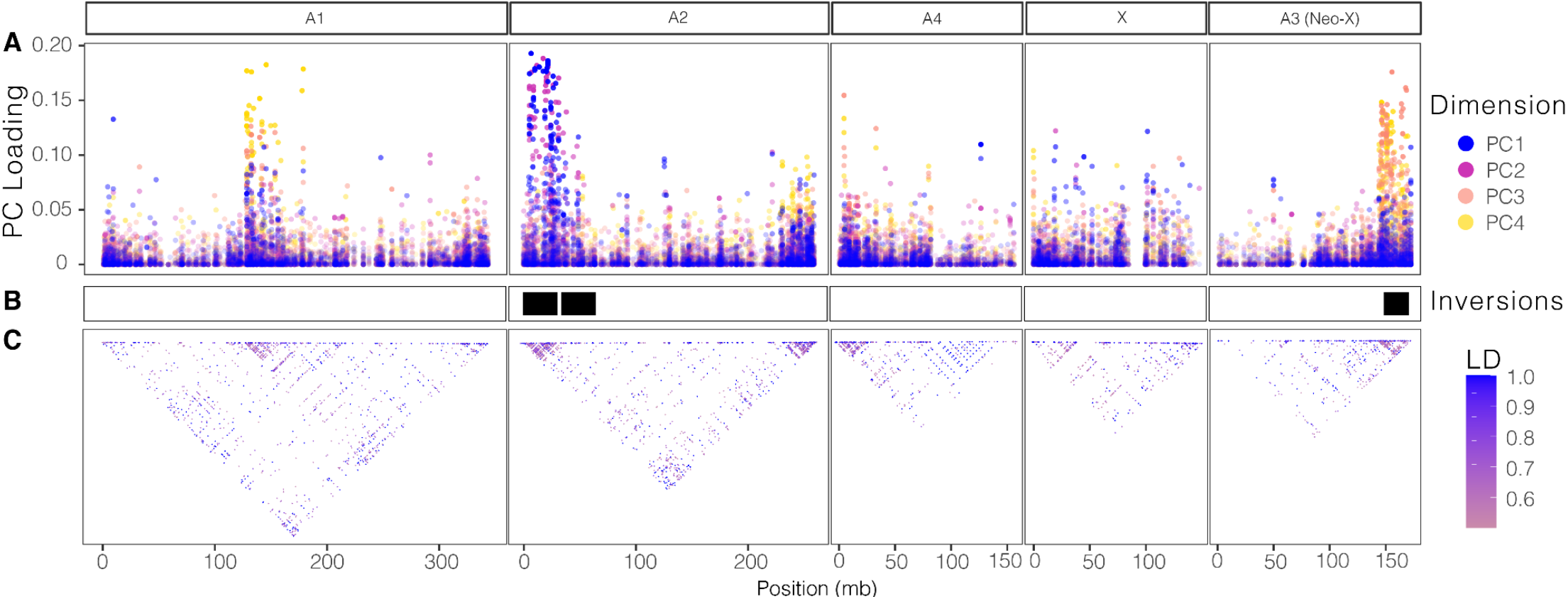
Genomic localization of SNPs participating in XX/XY-cytotype population substructure in *Rumex hastatulus*. **A**. Distribution of SNP effect sizes for XX/XY-cytotype PC dimensions 1-4 (color) along the chromosomes. **B**. Inversions between cytotypes shown as black blocks (data from Rifkin et al. 2020). **C**. Linkage disequilibrium greater than 0.5 (color) within XX/XY.

## DISCUSSION

Genomic analysis of the *Rumex hastatulus* hybrid zone offers rare insights into the evolution of reproductive isolation involving neo-sex chromosomes. Because inter-cytotype hybrids show reduced fertility from controlled experimental crosses (Kasjaniuk et al. 2018) and previous work inferred an absence of gene flow between allopatric populations of each cytotype (Beaudry et al. 2019), we might have expected to find little or no hybridization in the zone of contact. Instead, consistent with cytogenetic studies by Smith (1969), we found evidence of extensive hybridization between the two *R. hastatulus* cytotypes in the zone of contact. We detected geographic variation in the amount of hybridization and in the extent of gene flow across the genome. These findings allowed us to analyse both the structure of the hybrid zone and the role of neo-sex chromosomes in reproductive isolation, as we discuss below.

### Structure of the hybrid zone

The geographic region in which we identified hybrids is roughly consistent with cytological evidence from the 1960s (Smith 1969). In contrast with Smith’s findings, we observed almost total inviability of individuals from populations closest to the west side of the Mississippi river. In the glasshouse, none of the plants that originated from MLRA O (*Mississippi Delta Cotton and Feed Grains Region*) survived. Also, *R. hastatulus* individuals were absent from herbarium collections from the MLRA-O region (see Methods). The major characteristic of the MLRA-O area is a high-water table and excessive flooding (NRCS 2006), ecological conditions that are generally not suitable for the growth of *R. hastatulus* populations, which prefer drier soils (Conn and Blum 1981). Smith (1969) found signals of hybridization in this region whereas we only identified hybrids on the east side of the river. This leaves open the question as to whether individuals from populations in the MLRA-O region do not survive in the glasshouse because of unconditional hybrid inviabilities and/or because of an interaction with the growing conditions in the glasshouse. Although the question of whether hybrid individuals persist on the west side of the Mississippi river remains unclear, our study indicates that there is a strong signal of hybridization on the east side of the Mississippi river.

Patterns of hybridization are often used to infer the genetics of reproductive incompatibilities (Turelli and Moyle 2007; Turelli and Orr 2000). Indeed, the generation (*F*_1_, *F*_2_, etc) at which hybrids are inviable or infertile can be harnessed to understand the role of dominance in hybrid breakdown (Turelli and Orr 2000). Given the broad range of ancestry proportions and heterozygosity that we detected in our analyses, our results suggest most hybrid individuals appear to be late generation hybrids. Our findings showing strong evidence of a hybrid zone suggest that introgression is unlikely to be stalled at the *F*_1_ generation because of significant lethality or sterility, thus rejecting conclusions of a complete loss of gene flow between the cytotypes based on allopatric populations (see Beaudry et al. 2019). Similarly, asymmetries in the direction of gene flow are informative of the relative role of uniparentally inherited genetic factors (Turelli and Moyle 2007). By Bayesian genomic cline *(bgc)* analysis, we inferred a disproportionate effect of XX/XY alleles crossing the Mississippi river eastwards. The pattern we observed could be, in part, ecological, including for example influences of the direction of wind on pollen and seed dispersal, as seen in a recent meta-analysis of genetic diversity in trees (Kling and Ackerly 2021). However, previous experimental work suggests a role for genetics, as inter-cytotype crosses also suggest correspondingly asymmetric barriers to gene flow, which have parent-of-origin effects on hybrid male genotypes (Kasjaniuk et al. 2018). Overall, the characteristics of hybrids in the hybrid zone is suggestive of a heterogenous pool of hybrids, but is not directly informative about the genetic architecture of reproductive isolation.

When fitting clines to whole-genome estimates of hybrid index, we found that different estimates of hybridization returned different cline fits (Table 1). The different best-fitting models likely reflect differences between the assumptions of CONSTRUCT versus the (assumption-free) results of PCA. If we assumed the discrete classification of individuals into *K* ancestral groups, cline fitting returned a model supporting less gene flow across the hybrid zone. In contrast, fitting PC1 suggested a continuous distribution of genotype space across the entire range, including some shared polymorphism between allopatric populations as reflected in non-zero values of *p*. Models 2 and 4 were not significantly different in both hybrid index estimates, reflecting the challenges of distinguishing low levels of gene flow from ancestral polymorphism (Kern and Hey, 2017). However, the estimated effective migration surface (EEMS) makes few assumptions about discrete populations, as the number of demes increases, and indicated reduced gene flow in the hybrid zone, suggesting that gene flow is indeed reduced. Together, these results paint a broadly consistent picture of ongoing but limited hybridization with barriers to gene flow.

Empirical studies of dioecious plant species indicate that gene flow is often low in hybrid zones (Pickup et al. 2019). In *Silene latifolia and S. dioica* hybrid zones in Switzerland, differentiation genome-wide was low with late generation hybrids common but early generation hybrids mostly absent, suggesting historical introgression without extensive recent hybridization (Minder et al. 2007). The difference between the *Silene* and *Rumex* results may arise because *S. latifolia* and *S. dioica* are much more divergent in reproductive biology, phenology and habitat preferences (Baker 1948) than the *R. hastatulus* cytotypes we investigated here (Jackson, 1967; Puixeu et al. 2019). Minder et al. (2007) did not investigate the impact of the sex chromosomes of *S. latifolia and S. dioica*, as the study was based on unanchored amplified fragment length polymorphism (AFLPs). Given the rarity of analyses of hybrid zones in dioecious plants with sex chromosomes, it is clearly too early to make general conclusions on the nature of hybrid zones and their genomic consequences. Nonetheless, genomic heterogeneity in hybridization involving the sex chromosomes is common in animals (Hewitt 1975; Hooper et al. 2018; Macholan et al. 2011), and plants generally show variation in gene flow across the genome (Abbott 2017; Pickup et al. 2019) so it seems likely that overall patterns in plants will be similar to those in animals.

### Neo-sex chromosomes and reproductive isolation

Given the genomic evidence for the presence of ongoing introgression between cytotypes in the hybrid zone of *R. hastatulus*, we were particularly interested in assessing whether the sex chromosome and/or neo-sex chromosome showed evidence for reduced introgression. In scanning for the location of SNPs with large effects on genotypic differentiation between the cytotypes, we found that SNPs on the neo-X chromosome had disproportionately large loadings on the first principal component (PC1). This pattern is suggestive of an enrichment for loci with high differentiation between the cytotypes on the neo-X, which can indicate a role for the neo-X in reproductive isolation. Furthermore, when fitting clines SNP-by-SNP with *bgc*, we found that the small number of SNPs with a significantly steeper cline than the genome-wide average was on the X or neo-X chromosomes. In contrast, there was no signal of a disproportionate role of the neo-X in the genomic-regional clines: when fitting clines separately for each chromosome, split between regions of low and high recombination, the confidence intervals for cline widths mostly overlapped. This disagreement between the results for SNP-by-SNP clines and genomic-regional clines may be due to a lack of power to differentiate between genomic regions when fitting many cline parameters (six in our best fit models) to entire genomic regions; *bgc* avoids this need by significantly simplifying the model down to two parameters (Gompert and Buerkle 2011). Alternatively, these few outliers may be on the X and Neo-X themselves and their signal is overwhelmed by a chromosome-wide analysis. Indeed, the X chromosome of *R. hastatulus* is very large (∼20% of the 1.3GB genome) and therefore likely exhibits significant variation in gene flow among sites.

Our results are consistent with a role for the neo-sex chromosome in the evolution of reproductive isolation but leave the specific mechanisms unresolved. The appearance of late generation hybrids in our data suggests that reduced fertility from aneuploidy is unlikely to be the sole driver of reproductive isolation. Instead, the rearrangements could play a role in reproductive isolation by restricting recombination. Regions with high rates of recombination may be able to actively purge genetic incompatibilities, for example, as seen in swordfish hybrids (Schumer et al. 2018). However, while our data support an important role for neo-X linked SNPs in cytotype differentiation, we found no evidence that variation in recombination rate was involved.

Alternatively, theory shows that X-linked genes are more likely to contribute to local adaptation because of their relatively lower movement between populations of X chromosomes compared to autosomes (Lasne et al. 2017). In a large-scale geographical study of population differentiation in *R. hastatulus* involving the two cytoypes grown under uniform conditions in the glasshouse, Puixeu et al. (2019) demonstrated significant differentiation in sexually dimorphic traits associated with abiotic climatic variables, such as mean annual temperature and precipitation, suggesting local adaptation could be sex-specific. Linkage of sex chromosomes with alleles having locally sex-specific benefits would be consistent with the absence of hybrid breakdown in the contact zone and the lack of evidence for gene flow between allopatric populations (Beaudry et al. 2019). Although alleles can move between the cytotypes, albeit at a reduced rate, local adaptation may limit the success of migrating alleles across a broader geographic range. Mapping of quantitative trait loci for sexually differentiated traits in crosses between the cytotypes would be a promising approach to further understand the impact of sex-linkage and sex-specific local adaptation. Further disentangling the evolutionary forces that first led to the spread of the neo-X in populations in the eastern range of *R. hastatulus* may also provide valuable insights on the role of the neo-X in reproductive isolation.

### XX/XY substructure and chromosomal rearrangements

Unexpectedly, we observed three regions with disproportionately large contributions to population substructure within the XY cytotype, and our comparative mapping suggests two of these regions align with structural chromosomal rearrangements between the cytotypes. Large rearrangements appear to contribute to differentiation among geographic regions in many plant species (Rieseberg 2001). Often, such rearrangements are associated with differences in ecology and habitat differentiation, leading to the hypothesis that they play an important role in local adaptation (Connallon et al. 2018; Kirkpatrick and Barton 2006; Todesco et al. 2019; Yeaman 2013). Whether the rearrangements between the cytotypes of *R. hastatulus* are segregating within the XX/XY cytotype is unknown, but previous comparative mapping strongly suggests two of these rearrangements are inversions between the XX/XY and the XX/XY_1_Y_2_ cytotype chromosome order (Rifkin et al. 2020) and linkage disequilibrium data suggest lower recombination in these genomic regions. We did not observe any signals that the regions containing these inversions play a role in affecting rates of gene flow or in limiting hybridization between the cytotypes. Long-read whole genome resequencing could be used to investigate finer scale patterns of differentiation and perhaps capture segregating inversions. Investigating whether genetic variation within chromosomal regions containing structural variation is associated with ecological gradients across the geographical range of the XX/XY cytotype range should provide insight into the potential role of chromosomal rearrangements in population differentiation in *R. hastatulus*.

In conclusion, our findings indicate that despite large scale chromosomal rearrangements between the cytotypes of *R. hastatulus*, hybrids are viable and sufficiently fertile to participate in gene exchange over several generations in populations along a 200-300 km cline. Our results are consistent with a low but significant level of gene flow across the hybrid zone. We found a disproportionate role of the neo-X chromosome in restricting gene flow. Together, our results highlight the importance of genomic studies of hybrid zones to identify the role of neo-sex chromosomes in reproductive isolation in wild plants.

## ACKNOWLEDGEMENTS

The authors would like to thank Yunchen Gong for server help, Baharul Choudhury for wet lab assistance and Elsie Shogren for comments on the manuscript. This research was funded by Discovery Grants from the Natural Sciences and Engineering Research Council to SIW and SCHB. JLR was supported by an EEB post-doctoral fellowship by NSERC grants to SIW and SCHB and FB was supported by NSERC grants to SIW.

## Data Accessibility

All code developed and used in this script are available on github (github.com/felixbeaudry/rumexHybridZone). All new data (raw reads for 143 contact zone individuals) will be available for download on NCBI’s SRA upon publication.

## Author Contributions

SCHB, SIW, FEGB and JLR conceived and designed the study; FEGB and JLR performed the field sampling; FEGB, DK and MJ-C prepared the samples for sequencing, FEGB, JLR, AP and SIW conducted the analyses; and all authors contributed to the writing of the manuscript.

## SUPPLEMENTAL TABLES

**Table S1.** Mean and variance across populations of *Rumex hastatulus* in number of offspring surviving to first flower for each maternal parent grown in the glasshouse. Location of populations with respect to latitude, longitude, state and population’s side of the Mississippi river (river-side).

| Pop | Latitude | Longitude | State | River-side | Mean progeny per maternal parent | Variance in progeny among maternal families |
| --- | --- | --- | --- | --- | --- | --- |
| ALVA | 33.63 | -89.69 | MS | East | 0 | 0 |
| ARCA | 32.6 | -92.98 | LA | West | 6.9 | 1.19 |
| BIEN | 32.35 | -92.9 | LA | West | 2.5 | 1.19 |
| BRAN | 32.16 | -89.88 | MS | East | 0.2 | 0.13 |
| BUCK | 33.33 | -93.47 | AR | West | 0.4 | 0.16 |
| CALH | 32.5 | -92.41 | LA | West | 2.6 | 0.86 |
| CALV | 31.96 | -92.79 | LA | West | 0.4 | 0.22 |
| CHUN | 32.31 | -88.88 | MS | East | 0 | 0 |
| CROS | 33.12 | -91.89 | AR | West | 0.5 | 0.22 |
| ELDO | 33.14 | -92.59 | AR | West | 2 | 0.56 |
| ENON | 31.26 | -89.67 | MS | East | 2.4 | 1.3 |
| FOUK | 33.29 | -93.84 | AR | West | 0.3 | 0.15 |
| GLEN | 31.11 | -92.64 | LA | West | 1 | 0.52 |
| HAMB | 33.29 | -91.69 | AR | West | 0 | 0 |
| HESS | 31.04 | -92.16 | LA | West | 0 | 0 |
| JONE | 31.5 | -93.56 | LA | West | 4.7 | 0.79 |
| KELL | 31.06 | -90.37 | MS | East | 0.7 | 0.3 |
| LEES | 31.2 | -93.12 | LA | West | 3.6 | 0.85 |
| LOUI | 32.18 | -89.22 | MS | East | 0.7 | 0.42 |
| MAGN | 33.25 | -93.22 | AR | West | 0 | 0 |
| MTHO | 33.26 | -92.96 | AR | West | 0.7 | 0.50 |
| NWHO | 31.21 | -89.88 | MS | East | 1.4 | 0.78 |
| OGAC | 31.04 | -88.97 | MS | East | 1.7 | 0.54 |
| PATT | 31.17 | -90.72 | MS | East | 0 | 0 |
| RUST | 32.52 | -92.64 | LA | West | 6.5 | 1.0 |
| SIBL | 32.58 | -93.3 | LA | West | 5.4 | 1.5 |
| STRO | 33.13 | -92.25 | AR | West | 1.7 | 0.90 |
| SYLV | 32.01 | -89.38 | MS | East | 2.3 | 0.80 |
| UTIC | 32.17 | -90.18 | MS | East | 1.1 | 0.41 |
| WEIR | 33.36 | -89.33 | MS | East | 0.3 | 0.15 |

**Table S2.** Summary of 14 models compared for cline fitting of hybrid index (HI) estimated from both principle component 1 (PC1) and CONSTRUCT for *Rumex hastatulus* cytotypes based on models from *HZAR*.

|  |  |  |  |  |
| --- | --- | --- | --- | --- |
| Hybridization Estimate | Model Number | Allele frequency (p) bounds | Cline function | $\delta$ and $\tau$ parameters |
| PC1 | 0 (Null) | N/A | N/A | N/A |
|  | 1 | [0:1] | logit | N/A |
| | 2 | [0:1] | $\delta$ and $\tau$ | mirrored |
| | 3 | [0:1] | $\delta$ and $\tau$ | independent |
|  | 4 | [p<sub>min</sub>:p<sub>max</sub>] | logit | N/A |
| | 5 | [p<sub>min</sub>:p<sub>max</sub>] | $\delta$ and $\tau$ | mirrored |
| | 6 | [p<sub>min</sub>:p<sub>max</sub>] | $\delta$ and $\tau$ | independent |
| CONSTRUCT HI | 0 | N/A | N/A | N/A |
|  | 1 | [0:1] | logit | N/A |
| | 2 | [0:1] | $\delta$ and $\tau$ | mirrored |
| | 3 | [0:1] | $\delta$ and $\tau$ | independent |
|  | 4 | [p<sub>min</sub>:p<sub>max</sub>] | logit | N/A |
| | 5 | [p<sub>min</sub>:p<sub>max</sub>] | $\delta$ and $\tau$ | mirrored |
| | 6 | [p<sub>min</sub>:p<sub>max</sub>] | $\delta$ and $\tau$ | independent |

**Table S3.** Results of cline fitting models with HZAR using Hybrid Index (HI) estimated at the population-level by CONSTRUCT for each chromosome of *Rumex hastatulus*.

| Chromosome | Model | AICc | Parameter | Estimate | Confidence interval |
| --- | --- | --- | --- | --- | --- |
| A1 | 4 | 20.48 | Center (Lon.) | -89.06 | [-89.47;-88.14] |
|  |  |  | Width (km) | 196.35 | [97.59;460.78] |
|  |  |  | p<sub>min</sub> | 0.024 | [1.64e-06;0.087] |
|  |  |  | p<sub>max</sub> | 0.99 | [0.95;1.00] |
| A2 | 2 | 21.75 | Center (Lon.) | -88.62 | [-89.33;-87.65] |
|  |  |  | Width (km) | 392.58 | [143.10;593.62] |
|  |  |  | δ (km) | 335.06 | [101.41;509.79] |
|  |  |  | τ | 0.0016 | [4.94e-06;0.33] |
| A3/Neo-X | 4 | 24.44 | Center (Lon.) | -88.97 | [-89.31;-88.19] |
|  |  |  | Width (km) | 155.89 | [75.95;389.87] |
|  |  |  | p<sub>min</sub> | 0.031 | [0.01; 0.07] |
|  |  |  | p<sub>max</sub> | 0.97 | [0.90;0.99] |
| A4 | 1 | 19.83 | Center (Lon.) | -88.51 | [-89.25;-87.58] |
|  |  |  | Width (km) | 459.27 | [332.72;633.71] |
| X | 4 | 21.03 | Center (Lon.) | -88.79 | [-89.32;-87.78] |
|  |  |  | Width (km) | 255.97 | [121.61;494.2] |
|  |  |  | p<sub>min</sub> | 0.022 | [0.00024;0.091] |
|  |  |  | p<sub>max</sub> | 0.99 | [0.95 ;1.00] |

## SUPPLEMENTAL FIGURES

**Figure S1.**
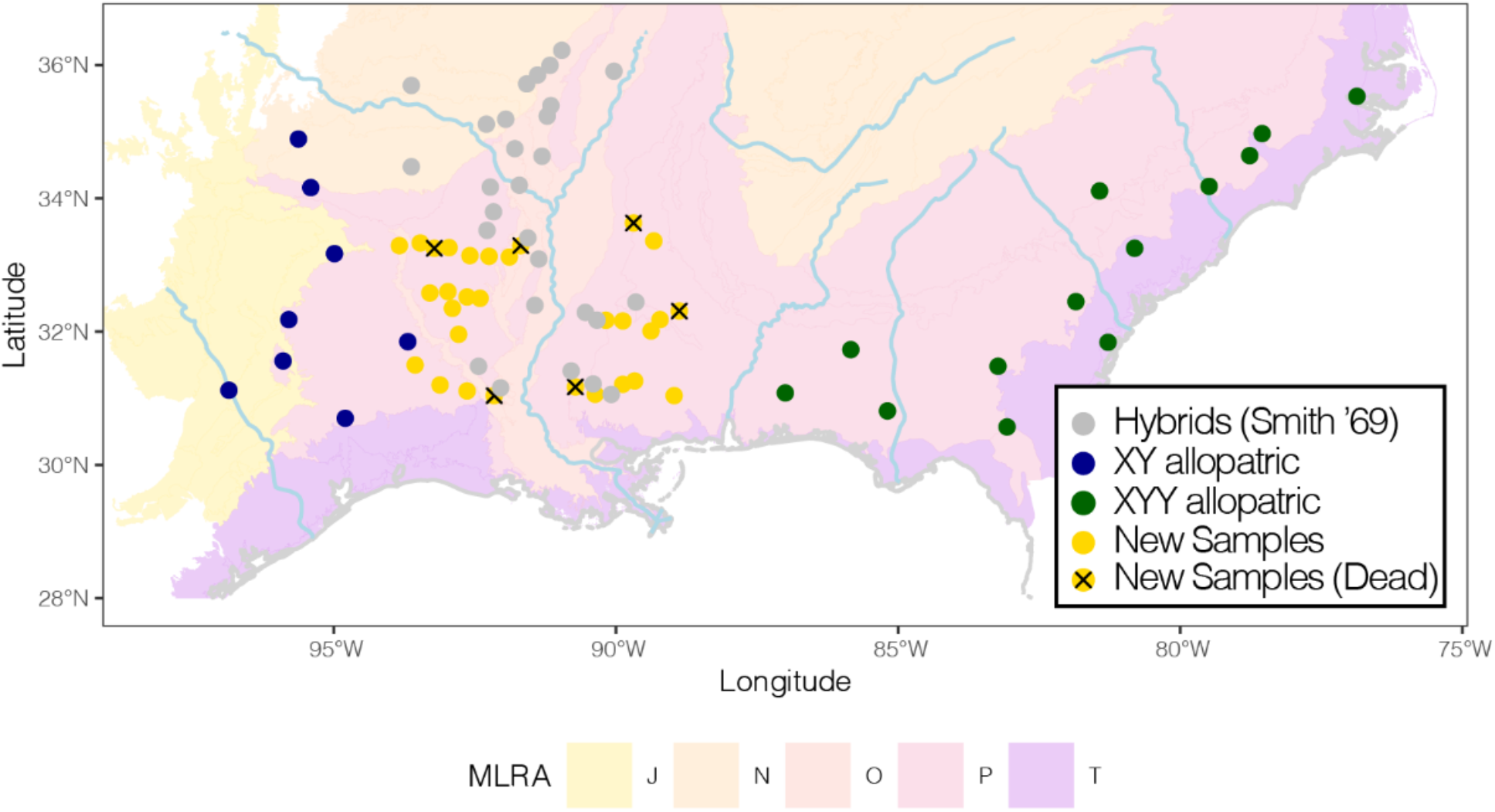
The geographical distribution of populations of *Rumex hastatulus* investigated in this study. Populations of the XX/XY- and XX/XY_1_Y_2_ cytotypes are in blue and green, respectively (allopatric in squares; sampled in Pickup et al. 2013, sequenced in Beaudry et al. 2019), and contact zone samples (collected for this study; circles) are in grey and successfully grown to first flower in black. Rivers in blue and state lines in grey. Background colors are Major Land Resource Areas (MLRA) as delineated by the Natural Resources Conservation Service.

**Figure S2.**
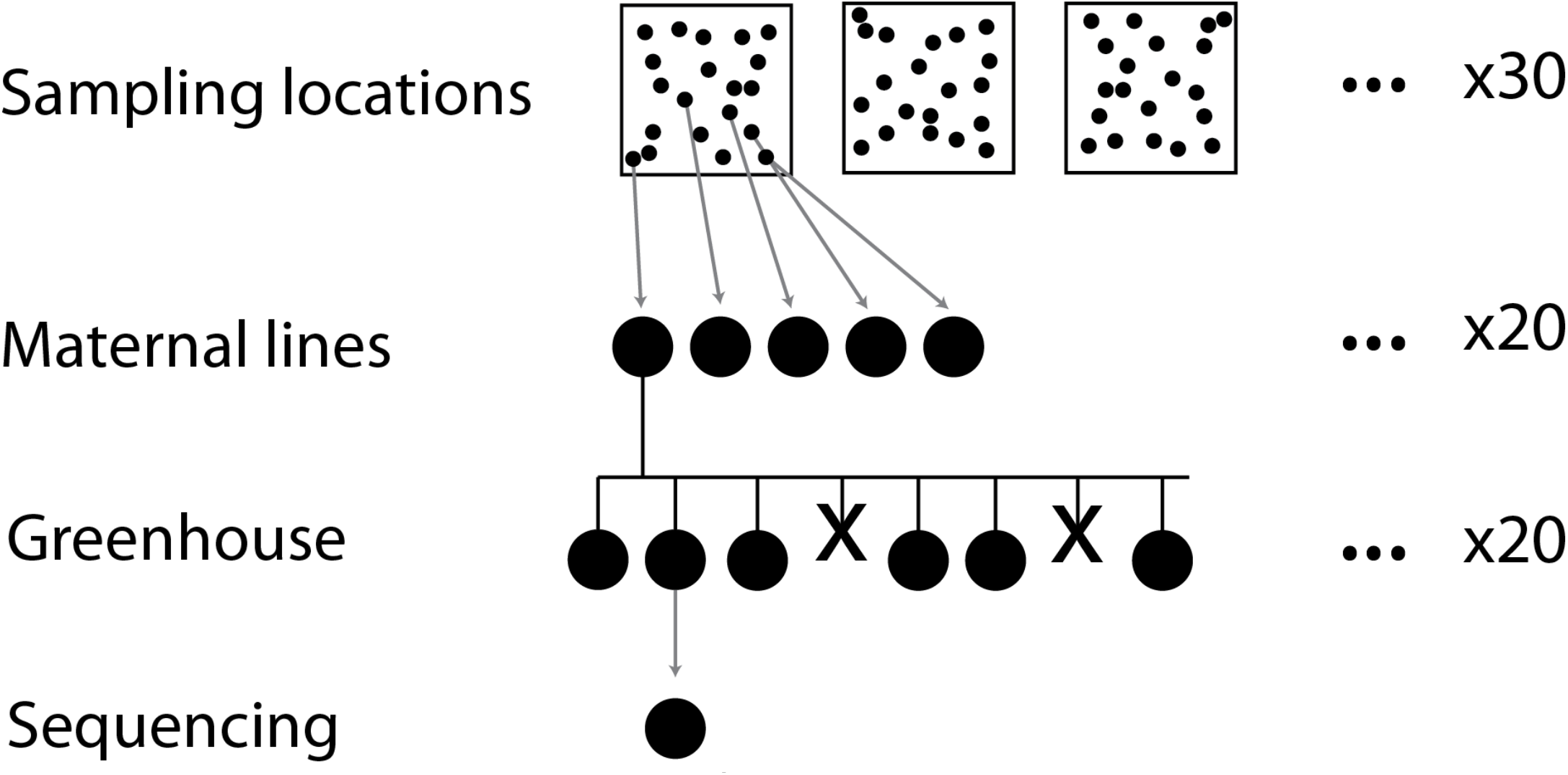
Sequencing design. At each of 30 sampling locations, we sampled seeds from a minimum of 20 open-pollinated females. For each sampling location, seeds from 20 females were chosen to start maternal lines. For each maternal line, we planted 20 seeds in the greenhouse. Not all individuals survived to first flower (represented with an ‘x’). For each maternal line, we sequenced one individual.

**Figure S3.**
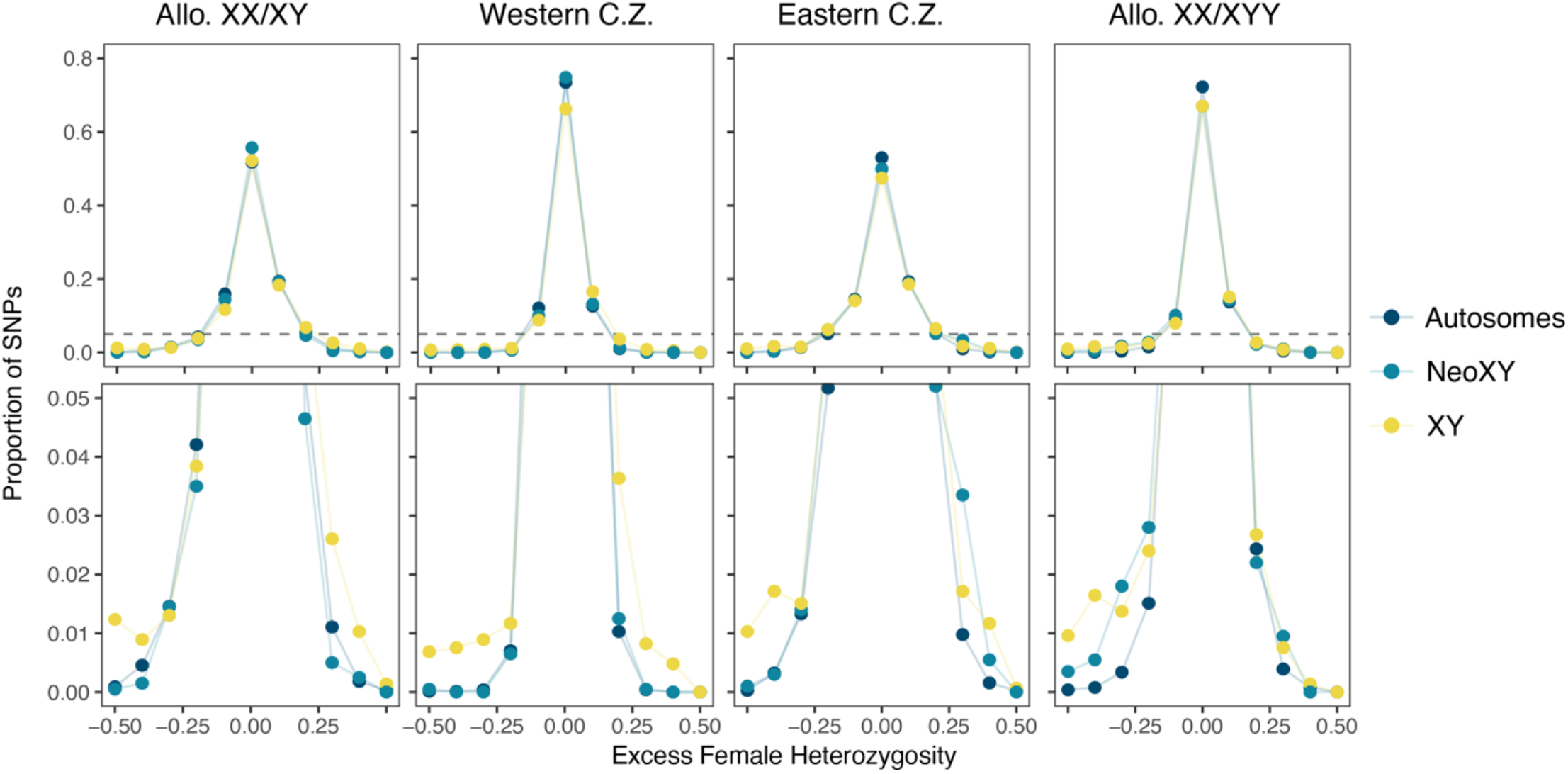
Heterozygosity (*2pq*) in females compared to males (excess female heterozygosity) in *Rumex hastatulus* populations at each of four locations (from west to east): allopatric XX/XY, western contact zone (C.Z.), eastern contact zone and allopatric XX/XY1Y2 for autosomes, neo-XY, and XY. Top row illustrates the full distribution, bottom panel is cut off at y=0.05 (dashed line in top row) to show differences in frequencies at SNPs with high levels of heterozygosity difference between males and females (both rows show the same data).

**Figure S4.**
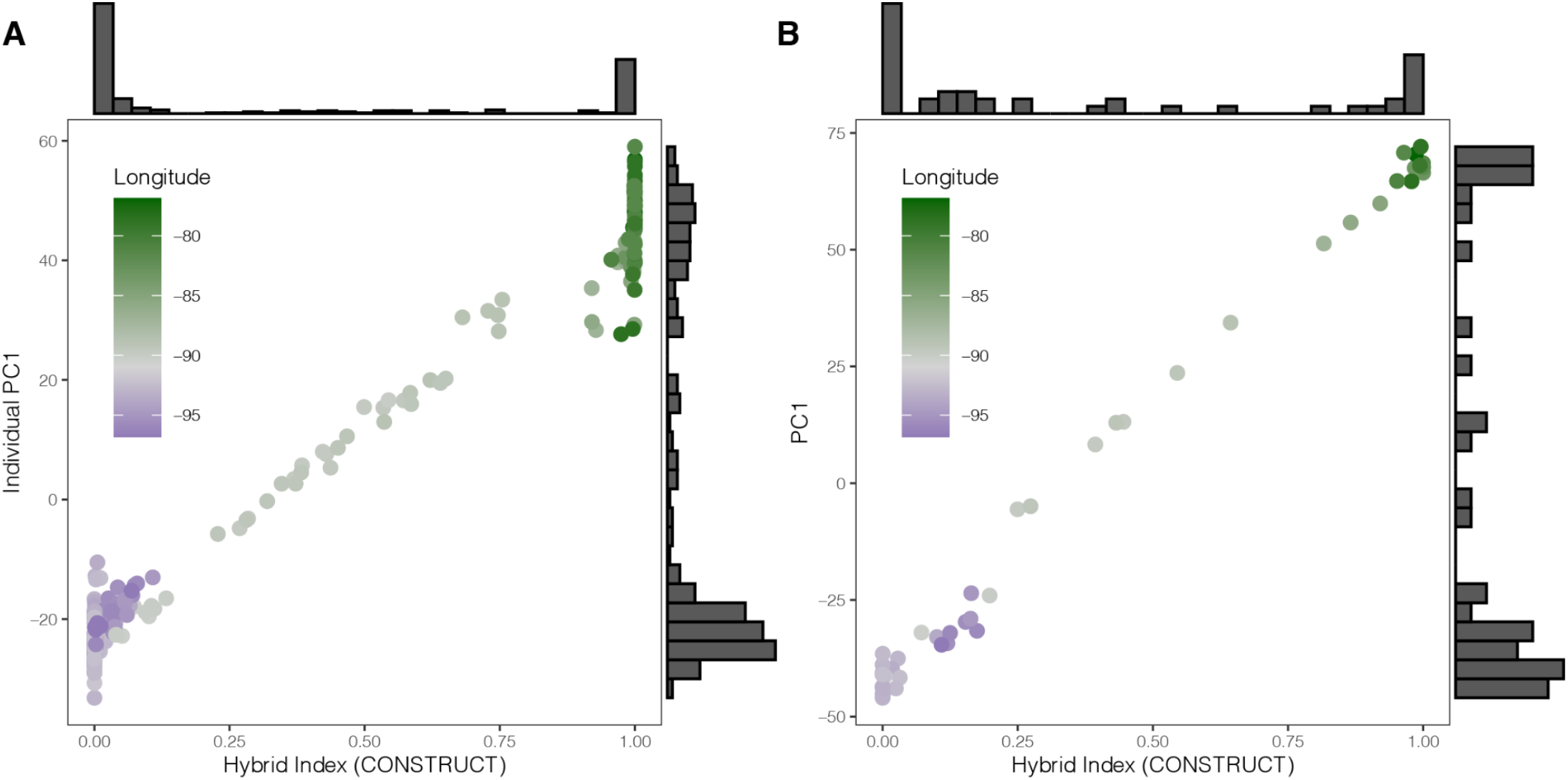
Relationship between PC1 and Hybrid Index estimated at the (**A)** individual-level (genotype-based) and **(B)** population-level (allele frequency) for populations of *Rumex hastatulus*.

**Figure S5.**
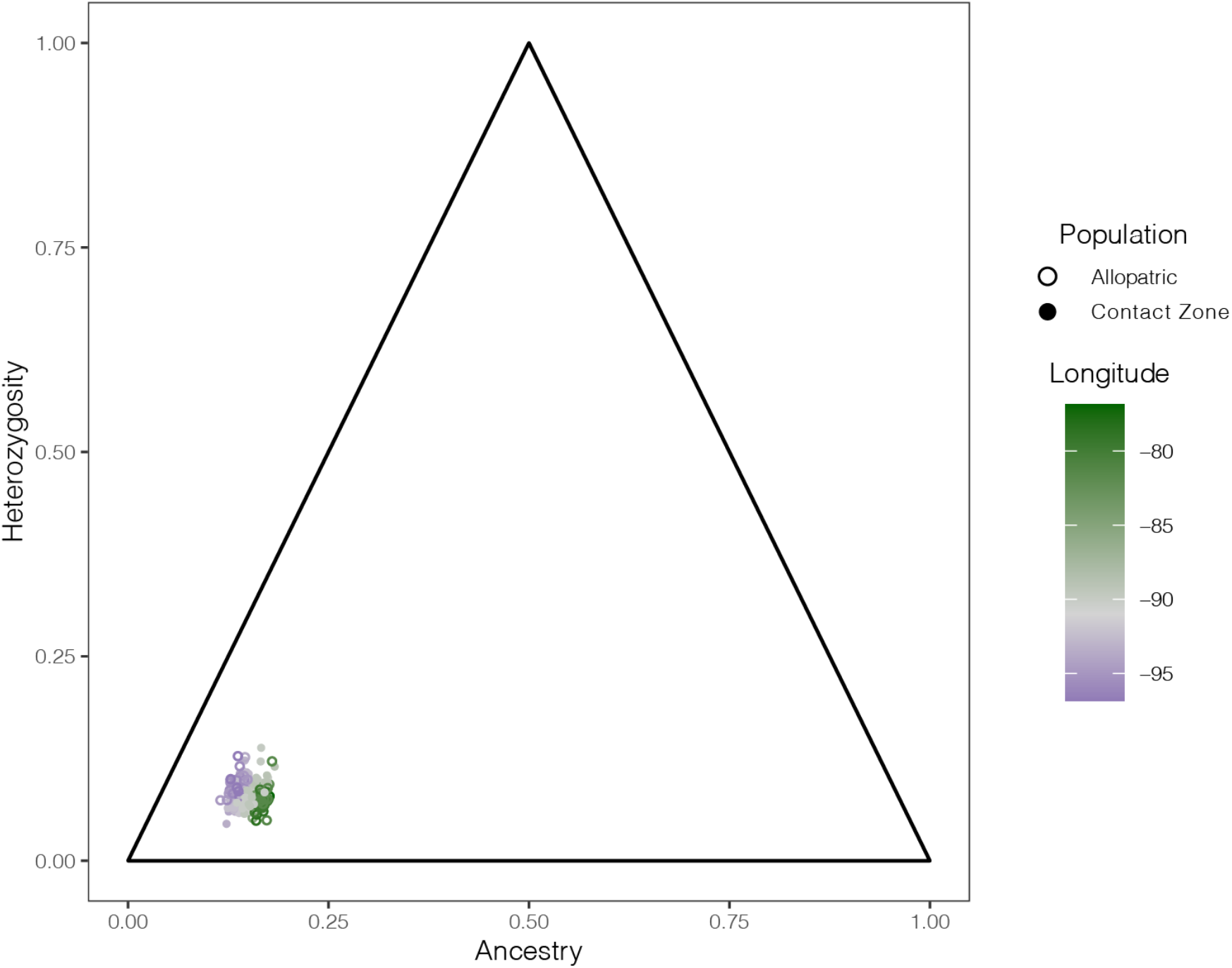
Inferring hybridization between cytotypes of *Rumex hastatulus*. Estimating hybrid generation breakdown genome-wide by assessing estimated ancestry compared to heterozygosity as estimated in HIest. Individuals are colored by longitude at which they were sampled and shapes represent whether the individuals were collected from a contact zone or allopatric population.

**Figure S6.**
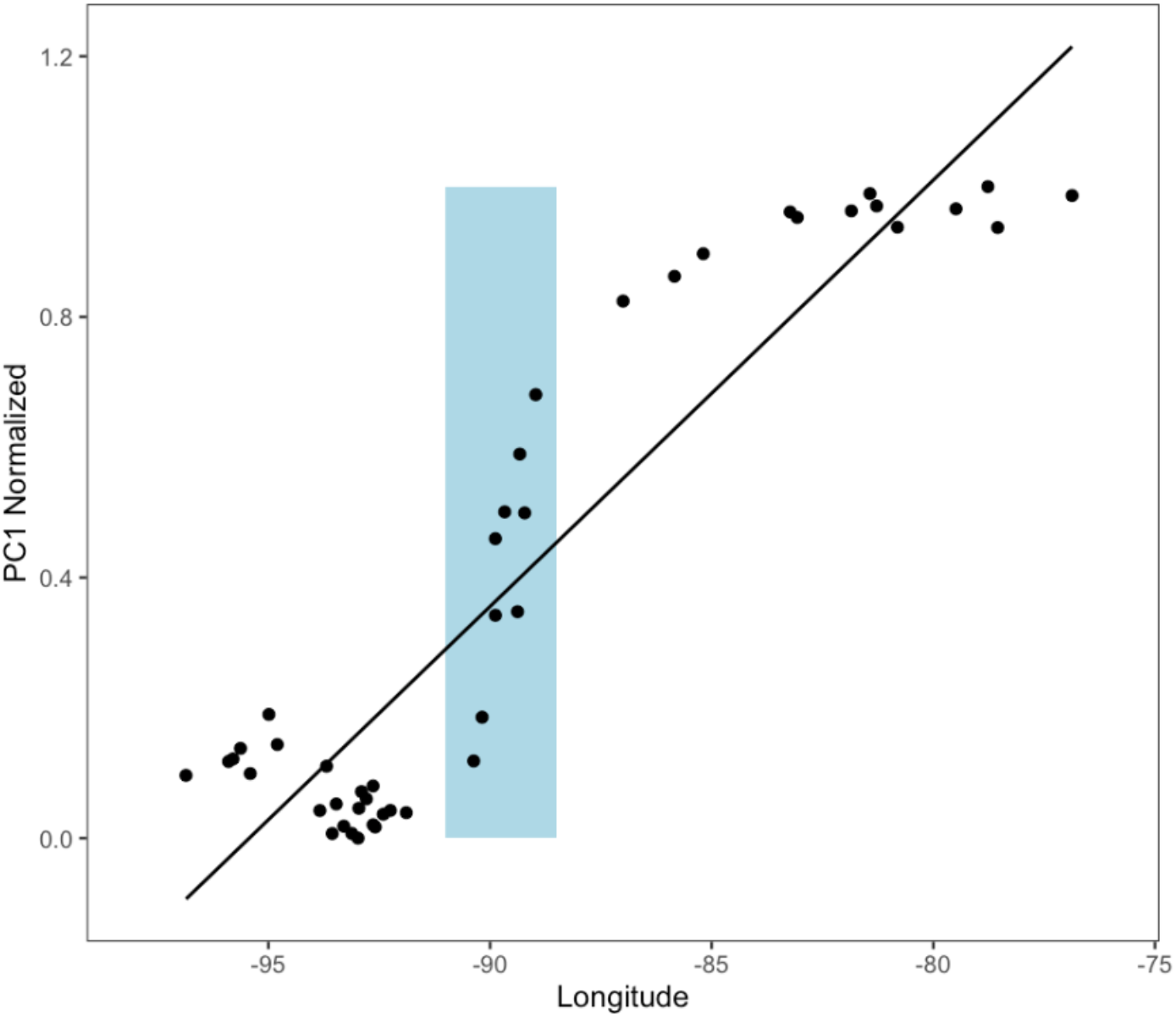
Relation between longitude and PC1 estimated at the population-level (using allele frequencies) in *Rumex hastatulus*. Line is linear model (y ∼ x + b) and blue square is hybrid zone.

**Figure S7.**
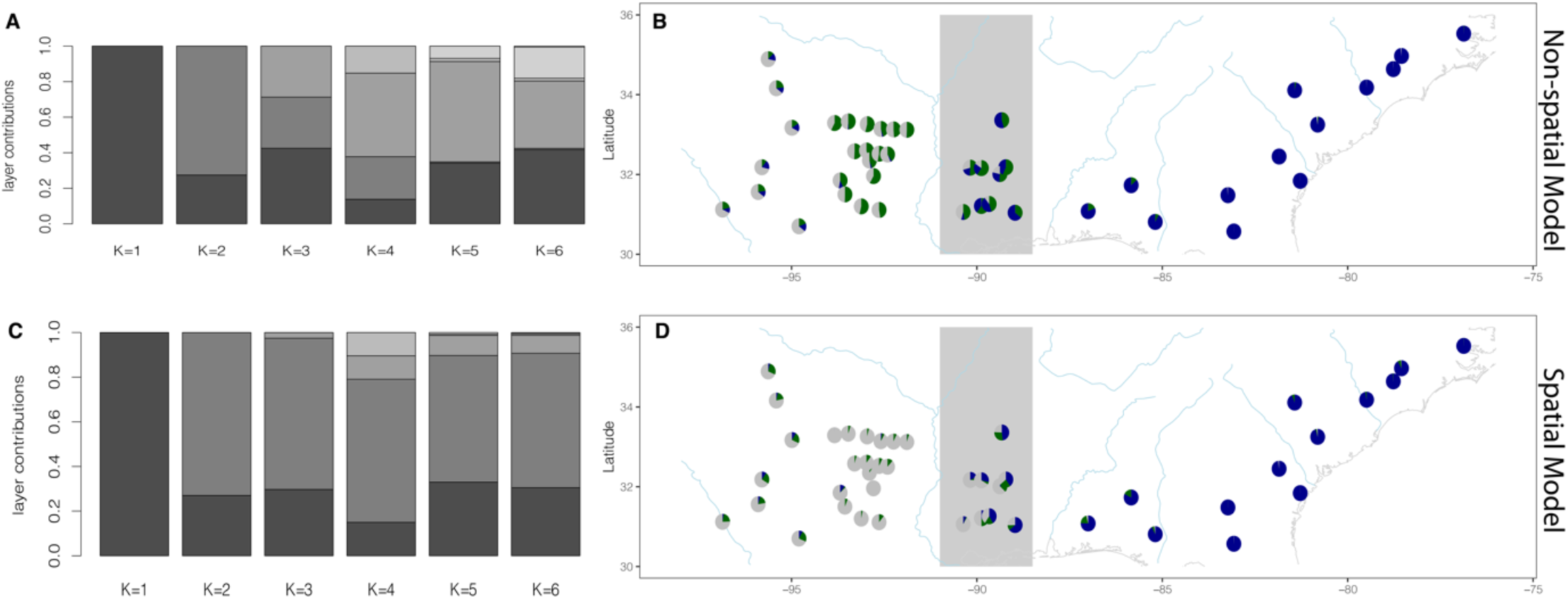
Comparison of spatial (bottom) and non-spatial (top) runs of CONSTRUCT based on SNP data from *Rumex hastatulus*. **A** and **C**. Variance explained by the contribution of each layer (shade of grey), with increasing value of K in *CONSTRUCT*. The comparison between the variance explained by increasing K is to diagnose the most informative value of K given the data. **B** and **D**. Pie charts for K= 1-3 (by color) under the assumptions K=3. Blue lines are rivers and the grey box is the hybrid zone.

**Fig S8.**
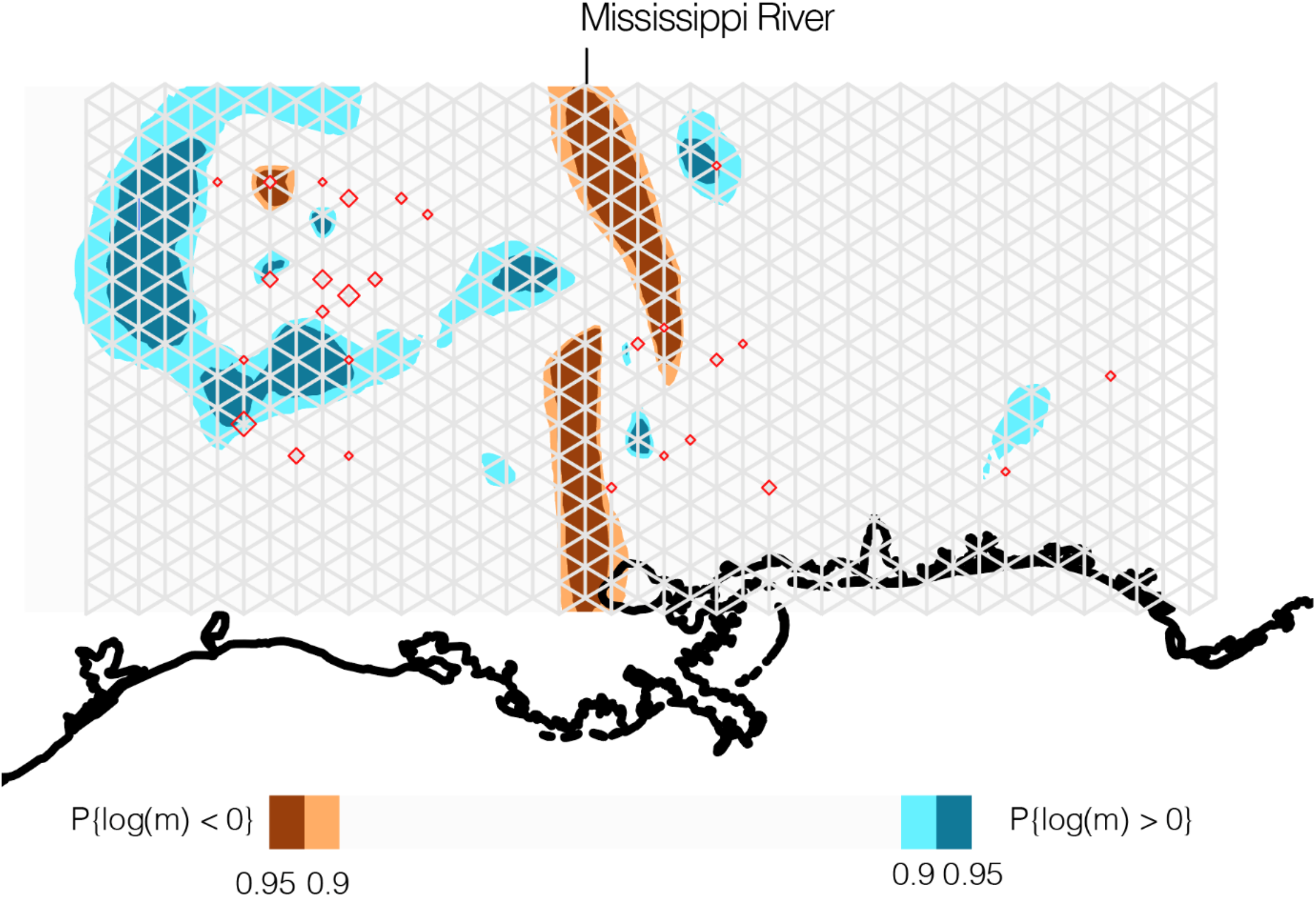
Posterior probabilities that the log migration rate is greater than or less than zero for the Estimated Effective Migration Surface (*EEMS*) run exclusively on contact zone populations of *Rumex hastatulus*. Red diamonds are the location of our population and diamond size is the relative number of samples from that location.

**Figure S9.**
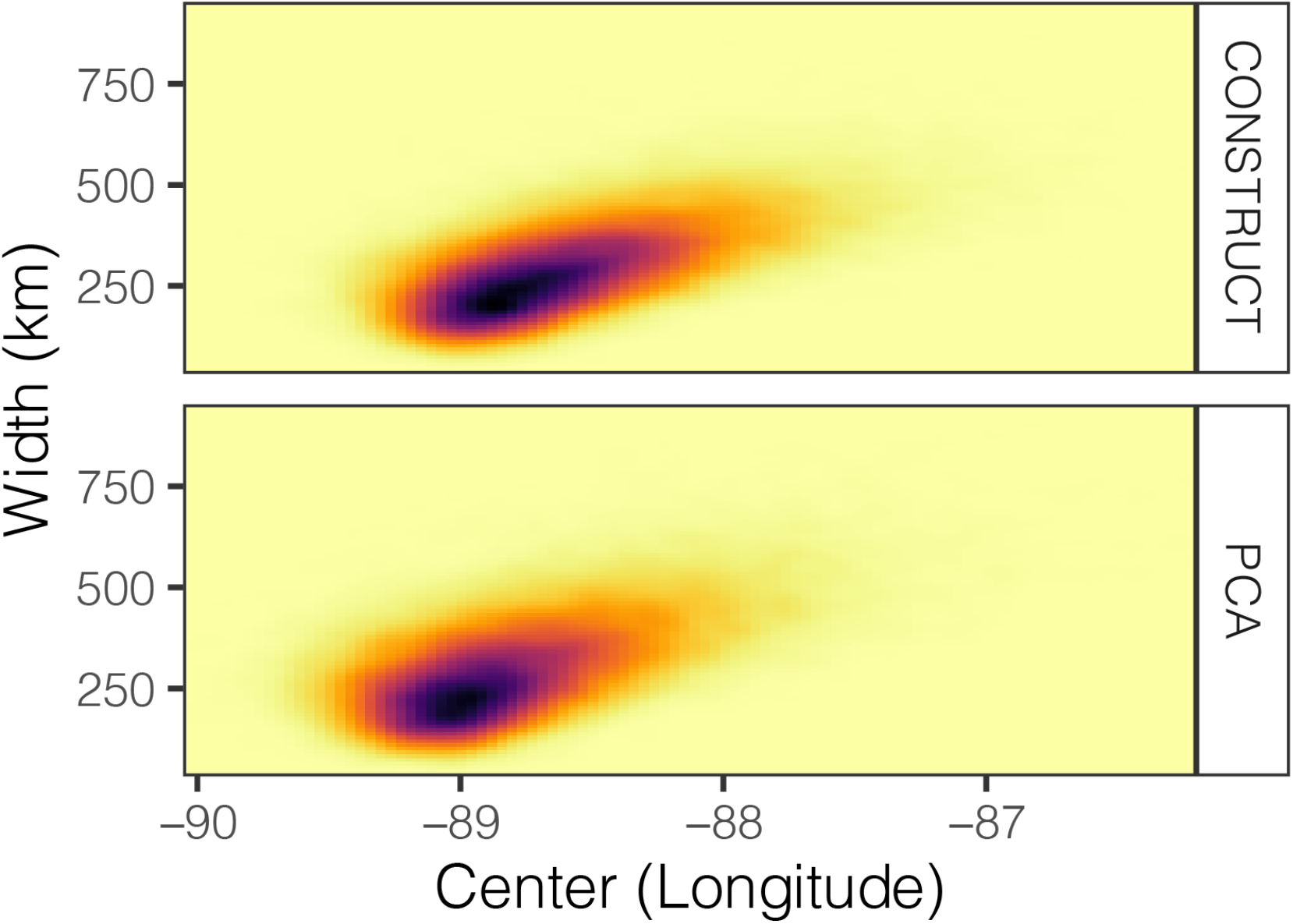
Relation between *Rumex hastatulus* cline center (in degrees longitude) and cline width (kms) for the cline fit to the CONSTRUCT data (top panel) and Principle Component Analysis (PCA; bottom panel) across the MCMC chain for HZAR. Darker blue represents more steps in the chain spent in that parameter area

**Figure S10.**
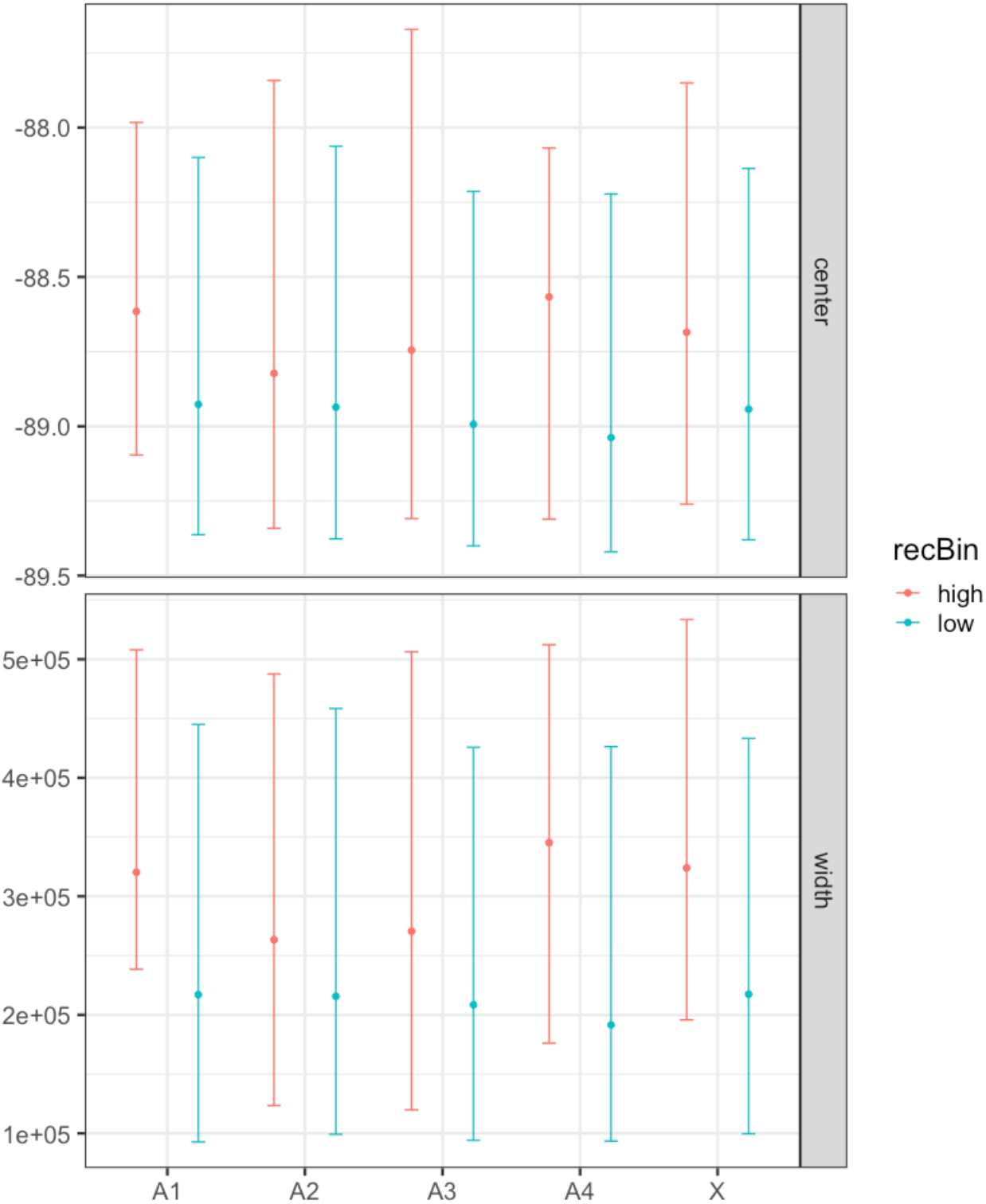
Estimates of cline center (longitude) and width (km) for each *Rumex hastatulus* chromosome according to bins of recombination (high and low recombination in red and blue, respectively). Error bars indicate the 95% confidence intervals.

